# ENVIROME-WIDE ASSOCIATIONS ENHANCE MULTI-YEAR GENOME-BASED PREDICTION OF HISTORICAL WHEAT BREEDING DATA

**DOI:** 10.1101/2022.08.14.503901

**Authors:** Germano Costa-Neto, Leonardo Crespo-Herrera, Nick Fradgley, Keith Gardner, Alison R. Bentley, Susanne Dreisigacker, Roberto Fritsche-Neto, Osval A. Montesinos-López, Jose Crossa

## Abstract

Linking high-throughput environmental data (enviromics) into genomic prediction (GP) is a cost-effective strategy for increasing selection intensity under genotype-by-environment interactions (G×E). This study developed a data-driven approach based on Environment-Phenotype Associations (EPA) aimed at recycling important G×E information from historical breeding data. EPA was developed in two applications: (1) scanning a secondary source of genetic variation, weighted from the shared reaction-norms of past-evaluated genotypes; (2) pinpointing weights of the similarity among trial-sites (locations), given the historical impact of each envirotyping data variable for a given site. Then, the EPA outcomes were integrated into multi-environment GP models through a new single-step GBLUP. The wheat trial data used included 36 locations, 8 years and 3 target populations of environments (TPE) in India. Four prediction scenarios and 6 kernel-models within/across TPEs were tested. Our results suggest that the conventional GBLUP, without enviromic data or when omitting EPA, is inefficient in predicting the performance of wheat lines in future years. However, when EPA was introduced as an intermediary learning step to reduce the dimensionality of the G×E kernels while connecting phenotypic and environmental-wide variation, a significant enhancement of G×E prediction accuracy was evident. EPA revealed that the effect of seasonality makes strategies such as “covariable selection” unfeasible because G×E is year-germplasm specific. We propose that the EPA effectively serves as a “reinforcement learner” algorithm capable of uncovering the effect of seasonality over the reaction-norms, with the benefits of better forecasting the similarities between past and future trialing sites. EPA combines the benefits of dimensionality reduction while reducing the uncertainty of genotype-by-year predictions and increasing the resolution of GP for the genotype-specific level.

---

Traditionally, the importance of genotype by environment interactions (G×E) has been seen as problematic and as needing to be controlled in plant breeding. Because of that, a large number of computational tools and statistical models have been developed and used for studying G×E in diverse scenarios and contexts from the evolutionary point of view for advanced multi-environment trials (MET). Essentially, the goal of this G×E analytics approach is to enable plant breeders to assess the stability of target traits under this MET, while supporting the prediction of the performance of newly developed cultivars evaluated under a pool of environmental conditions – also known as the target population of environments (TPE) of breeding programs.

However, even though MET are useful for selecting stable and high yielding varieties (Crossa et al., 2004) for the environments included in the trialing network, they are less suitable for providing targeted recommendations for specific environments (e.g., site and year combinations) where no field trials were performed. There is a rich history of statistical methods that incorporate external covariables in models to explain and understand the main environmental causes of G×E. The traditional regression (or reaction-norm) models of Yates and Cochran (1938), Finlay and Wilkinson (1963) and Eberhart and Russell (1966) describe environments based on the mean performance of cultivars in the test environments. However, instead of regressing the mean grain yield of cultivar performance on the mean grain yield of the environments (environmental quality index), environmental covariates can be used to characterize both (1) the environments in the MET and also (2) new target environments (Hardwick and Wood, 1972) that were not included in the MET trials.

Fixed-effect linear-bilinear models, such as the Sites Regression (SREG) (Crossa and Cornelius, 1997) and the Additive Main effect and Multiplicative Interaction (AMMI) (Gauch, 1988; Cornelius et al., 1996), are generalizations of single regressions on environmental data. They are used for studying genotypic response patterns across environments. In these models, the response patterns of genotypes and environments can be visualized graphically using biplots that allow the breeder to observe the high performing genotype(s) in a region(s) and/or sub-region(s). Partial least squares (PLS) regression is also a type of fixed-effect linear model which allows external environmental and genotypic covariables to be directly incorporated into the model, and has been demonstrated to be useful for identifying the climatic causes of G×E or the genetic factors (e.g., molecular markers) influencing G×E (Aastveit and Martens, 1986; Helland, 1988; Vargas et al., 1998; 1999; Montesinos-Lopez et al., 2022). In fact, the first mention of the use of high-throughput environmental data (“enviromics”) in plant science involved the use of PLS for scanning mechanisms acting in the internal cell environment (Teixeira et al., 2011). This concept was recently popularized by some seminal papers such as Resende et al. (2021) and Costa-Neto et al. (2021b), who mostly addressed the benefits of using enviromics to aid genomic prediction based on prior studies involving the so-called reaction-norm GBLUP (Jarquin et al., 2014; Morais-Junior et al., 2018).

As modern plant breeding is an interdisciplinary field based on multi-dimensional data types, the importance of developing new enviromics pipelines is key for successful prediction of G× E (de los Campos et al., 2020; Crossa et al., 2021). In the field of genomics, for instance, the availability of high density, low-cost genetic markers has made it possible to saturate the genome with these markers and predict estimated breeding values (Bassi et al., 2015). This has increased the precision of genetic value prediction compared to that achieved with traditional pedigree information. Genomic data can also help assess chromosome regions, e.g., marker effects and patterns of (co)variability of marker effects linked to different environmental conditions. Since the analysis of genetic and genomic data is one of the most challenging statistical problems currently faced, different models from diverse areas of statistical research need to be integrated in order to make significant progress in understanding genetic effects and their interaction with environments. In addition, new environmental sensors and remote sensing technologies (Morisse et al., 2021) permit the real-time, dynamic collection of environmental covariables with very high resolution anywhere in the world. Thus, marker technology combined with the rapid digitalization of high density environmental covariables (EC) and in-season image capture offers a new set of opportunities for improving the performance of MET for greater targeting of plant breeding (Crossa et al., 2021).

Jarquín et al. (2014) and Heslot et al. (2014) introduced the concept of genome-based reaction norms to model G× E using many environmental covariates (e.g., weather data, soil properties, crop growth model outcomes). Since then, a new wave of studies focused on the prediction of unobserved genotypes in new untested environments (see Crossa et al., 2022). The so-called linear kernel-based reaction norm model of Jarquin et al. (2014) was utilized by Pérez-Rodríguez et al. (2015), employing pedigree and environmental covariates in cotton trials. Similar substantial gains in prediction accuracy of genotype performance were obtained by Cuevas et al. (2016; 2017; 2018; 2019), He et al. (2019) and Costa-Neto et al. (2021a) using non-linear genomic Gaussian kernel models for modeling G×E interaction over the conventional linear GBLUP kernel. However, to date, these studies have focused on genomic-enabled predictions but have neglected two important components: first, the uncertainty of predictions in new environments, which can be substantial, as demonstrated by de los Campos et al. (2020); and second, the detailed enviromics assessment needed for the quantification of soil and climatic variables. Both factors are necessary for studying G×E and for the prediction of genomic estimated breeding values under different environmental conditions.

To address this situation, we made some efforts in the past to develop public envirotyping pipelines, such as *EnvRtype* (Costa-Neto et al., 2021b), also tested the merit of nonlinear kernels for modeling the environmental similarities in genomic prediction (Costa-Neto et al., 2021a), and for designing new “environmental markers” to add value in the prediction platforms (Costa-Neto et al., 2021c) by increasing the resolution of the predictive models in reproducing the phenotypic plasticity of the plants, while supporting the optimization of training sets with less phenotyping effort. Other authors have also addressed this situation by including crop growth models as a supervised machine learning tool, deep learning approaches using long-term field trial data and using robust experimental trialing networks for training reaction-norm models using in-field sensors. Despite the benefits of all these approaches, the lack of future environmental data is still a bottleneck in predicting yet-to-be-seen genotype-by-year interactions (G×Y). For “proof-of-concept” studies, it is easy to simulate “a new year” because we already have the environmental data observed in the past; however, for real-world breeding programs, we don’t have any environmental covariate for a yet-to-be-seen year. Because of this, here we focused on developing new tools for connecting past and future envirotyping data in order to reduce the uncertainty of G×Y predictions while using enviromics to add value in the current cultivar testing pipelines.

Here, to achieve this goal, we return to an old idea of connecting environmental and phenotyping data to interpret G×E interactions, considering a particular season and germplasm pool. To facilitate the interpretation of the concept, hereinafter we call it “environmental-phenotype association” (EPA). The theoretical basis of the EPA modeling approach assumes that the genotypic response over the environmental variation, and consequently the emergent G×E interaction observed in the experimental network, is strictly related to the particular envirotype-to-phenotype dynamics observed for each genotype and its evaluated growing conditions. Thus, the core of the reaction-norm not only determines the main drivers of the G×E (Millet et al., 2019; Costa-Neto et al., 2020a; Porker et al., 2020; Heinemann et al., 2022), but also explains how the environmental conditions have shaped the phenotypic correlations among experimental sites. Because of this property, here we used EPA as an intermediary step to adapt the envirotyping data (covariables) into environmental weights of similarity, while also exploring particular genotype responses for those factors (that is, reaction-norms) as a secondary source of genetic variation to approach genomics and the observed G×E. This analysis was also interpreted as a “reinforcement learning algorithm” capable of recycling past EPA as a precursor for future growing conditions and phenotypic responses.

To test this hypothesis, we analyzed a real-world breeding program and its intrinsic complexities observed in experimental trials (e.g., unbalanced conditions, diverse sets of locations across years). Within the CIMMYT Global Wheat Breeding Program, recent efforts have been made to describe, measure, and analyze G×E in MET tested breeding materials across major wheat growing areas around the world. This has shown that performance in CIMMYT’s main breeding and testing location in Mexico is correlated with various international sites belonging to TPE (or mega-environments, in this case) that represent the world’s major wheat producing areas (Rajaram et al., 1994; Braun et al., 1996; Crespo-Herrera et al., 2020).

As a TPE can be defined as a group of production environments that can be utilized for breeding in future years and/or in variable growing seasons, it is expected that the G×E may result from relatively static and predictable variation—for example, in soil or other conditions across the field—and dynamic, unpredictable and often significant temporal variability—i.e., weather over different years. The TPE delineations are based on climate, soil and hydrological characteristics and can also include socioeconomic factors, such as the resource levels of farm households. There are also different ways to group trials and environments into a TPE. Data from environmental sensors and satellites can be used to develop stratified hierarchical cluster analyses of sites and thus identify homogeneous environments wherein line performances will be highly correlated (Ornellas et al., 2019; Crespo-Herrera et al., 2021). Here we also check the merit of EPA as a data-driven approach in identifying those mega-environments and pinpoint the “essential trialing sites” within each environmental group, that is, identify the number of locations that represents the maximum diversity of growing conditions for each TPE (or across TPEs).

## METHODS

### CIMMYT historical wheat data

We used grain yield (GY, Mg ha^-1^) as the reference trait in this study. Data were collected from crop cycles 2011–2018 of Elite Spring Wheat Yield Trials (ESWYT) nurseries carried out in India and based on a previous and extensive study done by Crespo-Herrera et al. (2021) that aimed to define TPEs for CIMMYT wheat breeding in India (see Supplementary Figure 1). Here, we consider 370 wheat lines and 130 environments, which involved 36 locations, and 3 TPEs: TPE 1, the optimally irrigated Northwestern Plain Zone; TPE 2, the optimally irrigated, heat-stressed Northeastern Plains Zone; and TPE 3, the drought-stressed Central-Peninsular Zone. Details about the data involved in the experimental analysis, phenotype correction criteria and genotyping are given in Crespo-Herrera et al. (2021). Due to the highly unbalanced conditions (different genotypes across years and years with different locations), this data set is ideal for testing the prediction of untested genotypes in yet-to-be-seen years in a real-world breeding program. Details about how this trialing complexity was explored in different cross-validation scenarios are presented in subsequent sections.

### Envirotyping protocol

Here we considered “location” as a certain trialing site (geographical location), while “environment unit”, or simply “environment”, was considered as a combination of certain location × year × management. Thus, considering a historical breeding data, a certain location experienced a diverse set of environments across the years, also receiving a diverse set of genotypes evaluated at those environments. We considered two ways of envirotyping: one focused on characterizing “environments” using information for a single season (Steps 1 and 2 in the protocol detailed below) while a second was dedicated to characterizing “locations”, using the information across seasons (Steps 1, 2 and 3 in the protocol). Consequently, this resulted in two different environmental similarity matrixes (ERM). For “environments”, we used the conventional terminology of **W**-matrix, with dimension of environments × envirotyping covariables. For “locations”, we introduce the use of a new ERM based on the seasonal-averaged effects (**S-**matrix for locations), with dimension of locations × seasonal envirotyping covariables. Below we describe the four steps of those protocols.

#### Step 1: Raw data collection

The raw environmental data (soil and meteorological variables, Supplementary Table 1) was collected using remote sensing data bases, such as NASA POWER and in-field evaluations of soil data, as done by Crespo-Herrera et al. (2021), where more details can be found. Additionally, we computed variables with a higher biological meaning, such as growing degree days (GDD), vapor pressure deficit (VPD, kPa/day), evapotranspiration (ET0), atmospheric water balance (P-ETP), and the effect of temperature stress on radiation use efficiency, FRUE = {0,1}, as described in the *EnvRtype* package in R (Costa-Neto et al., 2021). The cardinal temperatures for GDD and FRUE calculations were assumed the same for pre and post anthesis stages, with Tmin = 0, Topt = 27.7, and Tmax = 40, as given in Wang and Engel (1998). We also subdivided the crop lifetime in “time windows” denoting a generalization of the expected development stage according to the Zadock’s scale for wheat (Zadoks et al. 1974), and following heat units for defining the windows of each stage (Rawson and Macpherson, 2000): (Stage 0 - 3) sowing to emergence, then to double-ridge (260° C.day^-1^); (Stages 3 to 5) double-ridge to terminal spikelet (+150° C.day^-1^); (Stages 5 to 6) terminal spikelet to heading (+350° C.day^-1^); (Stages 6 to 7) heading to anthesis (+° C.day^-1^); (Stages 7 to 9) anthesis to grain-filling and maturity (+500° C.day^-1^), where the symbol “+” denotes the accumulation of the given heat units for the current stage in relation to the previous stage.

#### Step 2: Computing envirotyping covariables (EC) for each environment

The computation of EC for each environment was done by a combination of meteorological variable x time window x quantile. Only soil covariables were considered static for each environment (e.g., clay content); that is, with no temporal scale across the crop lifetime, where we assumed EC as the combination of soil variable x quantile. Finally, a total of 108 high-quality ECs for each one of the 130 environments (combinations of year and locations) were found, resulting in 14,040 entries of envirotyping information. The final **W**-matrix (130 × 108) was mean-centered and scaled, with *w*_k_∼*N*(0,1).

#### Step 3: Computing ECs for each location across seasons

For each location, let’s assume that for each year of yield testing (each season), a specific **W**-matrix was considered. This is a season-specific matrix of envirotyping data and may not be repeatable across the years, that is, this is “snapshot” of the possible growing conditions a certain location may face. Each season has a diverse **W**-matrix, in terms of locations and the occurrence of environmental factors; however, the core of **W**-matrices across the years can be used to solve this and provide a “prior expectation” for a given location. It is possible to implement this only when a long-term time-series of envirotypic and phenotypic data is available for a given location. Thus, by using historical environmental data, it is possible to calculate the distribution of each combination of EC by time window across the seasons (assumed here as different years). This was implemented in terms of quantiles (10%, 50%, and 90%) of each EC for a given location, plus each static factor. Finally, the data were mean-centered and scaled, with *s*_*n*_∼*N*(0,1). After removing missing values and duplicated columns, the result was put into a matrix (hereinafter called an **S**-matrix) of *n* = 294 variables for each *l* = 36 locations, with **S** (*l* × *n*).

### Learning EPA through Partial Least Squares (PLS)

For the environmental-phenotype associations (EPA), we adopted the non-linear iterative PLS regression as an approach (NIPALS) due to its popularity, effectiveness for diverse data types and simplicity (e.g., Vargas et al., 1999; Teixeira et al., 2011; Monteverde et al., 2019; Porker et al., 2020; Montesinos-Lopez et al., 2022). The EPA analysis was performed in two ways, hereinafter referred to as PLS 1 (univariate PLS, focused on each genotype at the time) and PLS 2 (multivariate PLS, focused on the environmental relatedness), as detailed jointly with the genomic prediction models in further sections (see Supplementary Figure 2). Additional details of the NIPALS algorithm can be found in Palermo et al. (2009) and Sanchez (2012), as well as in the Appendix section. For each model, we also computed the Variable Importance in Projection (VIP) score. The VIP score has been used as a variable selection method, and can be computed for each covariable and latent factor as:

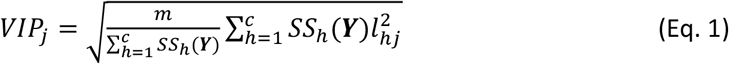

where ***Y*** is the matrix of responses, *m* is the number of variables in **X** (matrix of ECs, see Appendix Section); *c* is the number of components considered; 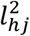 is the PLS weight of the *j*-th (j = 1, 2,…, *m*) variable for the *h*-th (h = 1, 2,…, *c*) at each component; and *SS*_*h*_(*Y*) is the proportion of **Y** explained by the *h*-th component. From the VIP, we calculated the relative VIP % by dividing each VIP by the maximum VIP for each genotype (PLS 1) or environment (PLS 2). The PLS approach was implemented in R using the *plsdepot* package (Sanchez, 2012).

### Integrating EPA outcomes in Predictive Models

Next, we introduced the use of EPA as an intermediary learning step in genomic prediction, using the multi-environment GBLUP model as a baseline. We used six models, in which the first three (M01-M03) were used as benchmark approaches and are currently used for multi-environment GBLUP, while the last three models (M04-M06) involve the inclusion of some EPA outcomes.

All models were fitted using the *EnvRtype*::kernel_model() functions, which is based on the kernel-optimized Hierarchical Bayesian approach implemented by the BGGE package (Granato et al., 2018). More details about this approach can be found at Costa-Neto et al. (2021a,b). We considered 15,000 iterations, in which the first 5,000 were used as burn-in considering a thinning of 5. The seed used was equal to *set*.*seed*(1112).

### M01: Conventional multi-environment GBLUP

Lopez-Cruz et al. (2015), Souza et al. (2017) and others, developed a conventional multi-environment GBLUP accounting for main genetic effects and kernel-based G× E. Here, we used this model as our benchmark and baseline structure, following:

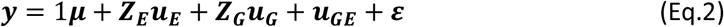

where ***y*** = [***y***_1_, …, ***y***_***n***_]’ are the vectors of observations collected in each of the *q* environments with *p* wheat lines; 1***μ*** is the vector of fixed intercept; ***Z***_***G***_ and ***Z***_***E***_ are the incidence matrix of genetic and environmental effects; ***u***_***E***_ is the vector of random environmental effects modeled by 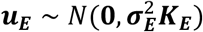, in which 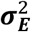 is the variance component related to the macro-environmental variation and ***K***_***E***_ is the kernel of macro-environmental effects, where the ERM were assumed as an identity matrix of *q* environments across the *p* genotypes ***K***_***E***_ = ***I***_***q***_ ⊗ ***J***_***p***_, that is, no prior relation was expected among environments. The genetic effects are modeled by a random main effect (***u***_***G***_), distributed as 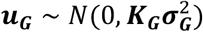, with ***K***_***G***_ = ***J***_***q***_ ⊗ ***G***, and a secondary kernel for environment-specific genetic effects due to G× E, represented by (***u***_***GE***_), with 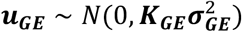 and modeled as multiplicative effects due to the GRM and ERM variations:

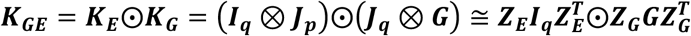

where 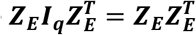, with ⊗and ⊙ denoted as the products of Kronecker’s and Hadamard’s, respectively. The matrix ***J***_***p***_ and ***J***_***q***_ denote the full-entry matrix of 1*s* with dimensions of *p* x *p* and *q* x *q*, respectively. Under balanced situations (every genotype at every environment), ***K***_***GE***_ could simply be computed as: ***K***_***GE***_ = ***I***_***q***_ ⊗ ***G***. Finally, the vector of residuals is assumed a relaxed form for residuals, with 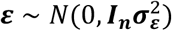.

### M02: Reaction-norm GBLUP with a linear kernel for W-matrix (*Ω*)

In this model (Jarquín et al., 2014) we considered the conventional use of envirotyping data (**W**-matrix), and consequently, also the linear variance-covariance matrix for environmental similarity as:

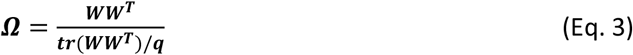

where ***W*** (*q* × *m*), with *m* environmental covariables (*m* = 108) and *q* environments. Consequently, the new kernel for modeling macro-environmental effects is now given by ***K***_***E***_ = ***Ω*** ⊗ ***J***_***p***_, and the subsequent G×E structure is:

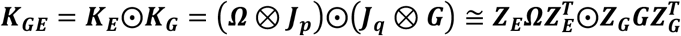

or simply ***K***_***GE***_ = ***Ω*** ⊗ ***G*** under balanced conditions as previously described. Thus, now the ***K***_***E***_ accounts for the degree of similarity among environments (given by a linear correlation), while the ***K***_***GE***_ is now the genotype-environment specific effects aimed to mimic reaction-norms.

### M03: Reaction-norm GBLUP with a nonlinear Gaussian kernel for W-matrix (*γ*)

From models M03 to M06, we adopted the Gaussian Kernel approach (GK) as the nonlinear method for modeling the environmental similarity (given by distances, He et al., 2019; Costa-Neto et al., 2021). In model M03, although the nonlinear ERM (***γ***) is computed using the same **W**-matrix in M02, we adopted a different notation to differentiate the linear kernel (***Ω***) and the nonlinear kernel (***γ***), which was estimated as:

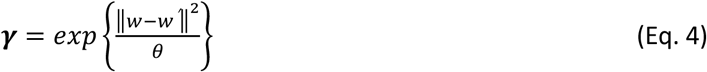

where ∥*w* − *w*’∥^2^ is the Euclidean Distance between each element of the **W**-matrix, and *θ* is a scaling factor assumed as the median value of the Euclidean distance matrix. Note that no bandwidth factor was considered in this equation (assumed as 1). Thus, model M03 is composed of ***K***_***G***_ = ***J***_***q***_ ⊗ ***G***, ***K***_***E***_ = (***γ*** ⊗ ***J***_***p***_) and 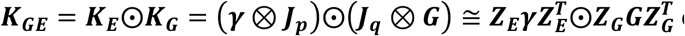 or simply ***K***_***GE***_ = ***γ*** ⊗ ***G*** under balanced conditions as previously described.

### M04: Reaction-norm GBLUP with environmental weights (*Φ*) from EPA

Here we introduce the EPA algorithm as an intermediary step in the multi-environment GBLUP. EPA is expected to be capable of linking historical phenotypic and envirotyping data, reducing the issue of lacking future envirotyping data while estimating the actual magnitude and impact of the environmental factors in the phenotypic variation observed in the trialing network. Below, we describe the four steps for incorporating EPA outcomes from the EPA algorithm in the baseline multi-environment GBLUP.

#### Step 1: Computation of the long-term envirotyping-based S-matrix

Now we consider the **S**-matrix instead of the **W**-matrix. As described in the Envirotyping Protocol section, the **S** is a matrix with dimension of location x seasonal ECs, where the seasonal ECs are a combination of ECs by quantiles (10%, 50% and 90%) across seasons; thus 108 × 3 = 324 seasonal ECs.

#### Step 2: Computation of the empirical Φ_0_ kernel of phenotypic correlations

To address the lack of information for future envirotyping data, this step aims to connect the past “envirotyping-realized” similarity into an “actual” similarity observed among environments, that is, the real phenotypic correlations among trialing sites. To implement this, the second step of this approach now focuses on computing a core of “empirical index” for a given trait, similarly as proposed by the conventional mean-centered average yield value approaches (Finlay and Wilkinson, 1963; Eberhart and Russell, 1966).

For each location, we computed quantiles of 10%, 50% and 90% for grain yield using all phenotypic records across years and genotypes. Then, we extracted a pool of “empirical environmental indices” (hereafter called an **S0**-matrix), which are based on empirical phenotypic data. Thus, while **S** contains the seasonal envirotyping data, the **S0** contains the seasonal descriptors of the environmental quality derived from the past observed phenotypes. This **S0**-matrix was then used to create an empirical location-relatedness kernel (***Φ***_0_, with *r* × *r*, where *r* is the number of locations), also following the nonlinear Gaussian kernel approach as:

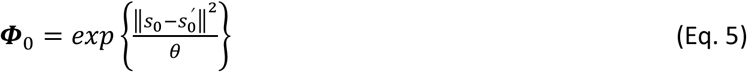

Note that this approach can be used considering an expected cropping time for a given location, which means that a statistic for each trialing site could accommodate a time-scale variation for diverse planting dates. Thus, for example, a same location could accommodate several “expected environments” by a combination of planting dates at each location. However, for the purpose of simplicity, and because most plant breeding programs perform analysis in almost the same planting seasons across years, here we consider the seasonal variations of envirotyping data for a static location and planting date.

#### Step 3: Translating envirotyping data into environmental weights (φ)

Next, the phenotype-based matrix ***Φ***_0_ is then dissected using the envirotyping-based **S**-matrix with the multivariate PLS enabled by the NIPALS algorithm (PLS 2, see Appendix). This approach can be mathematically represented as:

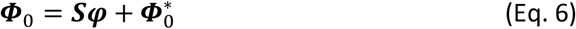

where ***Φ***_0_ (*r* × *r*) is an empirical (phenotype-based) relatedness; ***S*** (*r* × *n*) is the matrix of envirotyping data by location averaged across seasons; ***φ*** (*r* × *n*) is a matrix of estimated orthogonal PLS coefficients for each *n* envirotyping data, interpreted here as the *environmental weights* derived from the environmental-phenotype association and specific for each location; and 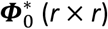 is the residual location diversity not captured by the envirotyping covariables in the **W**-matrix.

An interesting output of this analysis is the interpretation of the environmental factors that drive the similarity among locations. For this, the locations can be grouped using some clustering analysis (e.g., hierarchical clustering based on the principal components analysis). Finally, the interpretation of the VIPs scores (see Eq. 1) for each environmental covariable is a sign of how these factors have historically shaped the diversity among locations for a given trait in the breeding program.

#### *Step 4: Nonlinear kernel* (*Φ*) *using the environmental weights* (*φ*)

The environmental weights should be a better descriptor of the environmental similarities because they highlight how the growing conditions have affected phenotypic variation and experimental quality. Consequently, we propose using them as *markers of the environmental relatedness*, which can replace the conventional direct use of envirotyping covariables as done in the previous ERM approaches. Here, by using the nonlinear Gaussian kernel over the ***φ*** matrix, now it is possible to compute an environmental-similarity matrix for locations as:

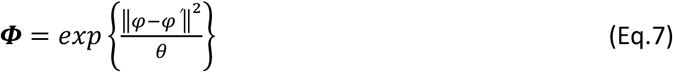

with ***Φ*** (*r* × *r*), thus, ***K***_***E***_ = (***Φ*** ⊗ ***J***_***p***_) and, consequently, 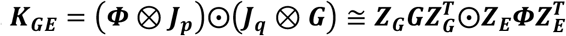.

### M05: Reaction-norm GBLUP with genotype-specific factors (R-matrix) from EPA

Model M05 introduces the use of a second EPA outcome. Here the focus is on studying the reaction-norm patterns from the genotypes in historical breeding trials, to finally recycling it as secondary source of genetic variation (hereinafter called **R**-matrix). This model is an update of model M04, that is, considering the protocols previously described for computing the ***Φ*** matrix.

#### *Step 1: EPA analysis for computing genotype-specific coefficients* (*λ*)

A site-regression (SREG, Crossa and Cornelius, 1997) model is fitted for each genotype with both phenotypic and envirotypic data available, which means that a specific regression model is fitted according to the number of observations available for a given genotype. The purpose of this approach is to dissect the genetic-plus G× E effects in terms of genotype-specific coefficients of reaction norm (hereafter named ***λ***) using the univariate PLS (PLS 1, see the Appendix). This modified SREG model using envirotyping data (Costa-Neto et al., 2020) is given by:

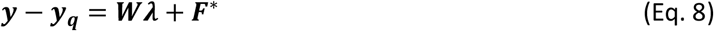

where ***y*** − ***y***_***q***_ (*q* × *1*) is a vector of phenotypic values for each genotype centered for each test environment; ***W*** (*q* × *m*) is a matrix of environmental covariables per environment (conventional envirotyping data); ***λ*** (*m* × *1*) is a vector of genotype-specific coefficients, assumed as the empirical reaction-norms for the given environmental covariable (reaction-norm); and ***F***^*^ is a vector of the residual SREG analysis not captured by the univariate PLS algorithm. Finally, after running this model for each *p* genotype, every vector containing the *m* genotype-specific coefficients was combined into a single matrix as ***Λ*** = [***λ***_1_, ***λ***_***2***_, ***λ***_***3***_, …, ***λ***_***p***_]***T***, with ***Λ*** (*p* × *m*). Due to the process of scanning reaction-norms into the modified SREG model, it is also possible to use the outcomes of the PLS approach to identify what environmental factors that most affect the performance of a certain genotype. The interpretation is done by analyzing the magnitude and direction (positive/negative) of the genotype-specific reaction-norms, while the variable importance on projection (VIP) could be used to score its importance. Because of this, now the EPA analysis integrates the classical G× E dissection studies (e.g., Crossa et al., 1999; Porker et al., 2019; Costa-Neto et al., 2020) into the current multi-environment genomic-enabled prediction platforms.

#### Step 2: Estimation of the adaptability through a reaction-norm based kernel (R)

At this step, we recycle the past reaction-norms as markers of the major G×E variation observed in the historical field trials. Then, the ***Λ*** matrix is used as genetic covariable, aimed to serve as a secondary source of environmental-specific genetic variation due to the shared reaction-norm patterns of the past evaluated genotypes. Because of the process carried out, this source of genetic variation does not follow an infinitesimal model and presents the same issues that are expected for environmental data (lack of linearity and additivity). Next, we used a nonlinear kernel (Gaussian Kernel) to translate the ***Λ*** matrix into a reaction-norm similarity matrix (***R***) as:

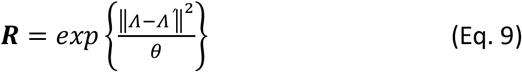

with ***R*** (*p* × *p*) if all genotypes are considered.

#### Step 3: Estimation of the reaction-norm weighted G-matrix (H)

In practice, plant breeders don’t have enough phenotypic information to compute the ***R***-matrix, especially if the goal is to predict the performance of a new genotype into a new season. To address it, here we propose to merge this ***R***-matrix with the conventional ***G***-matrix, in analogy of what is commonly done for combining pedigree and genomics in the so-called “single-step genomic prediction”. However, for this scenario, the ***G***-matrix (*p* × *p*) is known for all genotyped individuals (whether or not it was previously evaluated), but the **R**-matrix (*t* × *t*) only considers the empirical reaction-norm estimated for tested genotypes, that is, the training set, with *t* genotypes, in which *p* = *t* + *v*, and *v* is the number of yet-to-be tested genotypes (testing set). Because of this, by using the equation from Martini et al. (2018) it is possible to merge ***G*** and ***R*** into a so-called “**G**-matrix weighted by reaction-norms” (hereinafter called ***H*** or **H**-matrix), or simply a Single-Step Reaction-Norm model, computed as:

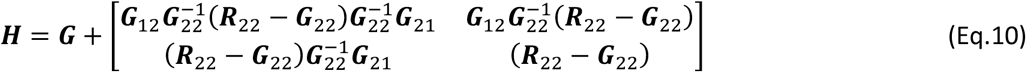

with

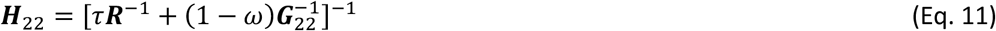

where ***H*** (*p* × *p*), *τ* and *ω* are scaling factors aimed at reducing inflations of the predictions and ensuring the convergence of the iterative approaches in the mixed models. To simplify the methodology, in this study the scaling factors were assumed as 1; however, it can be fine-tuned in further studies considering particular characteristics of each germplasm and trialing network.

Finally, the kernel-based G×E term was replaced from ***K***_***GE***_ to ***K***_***H***_ as 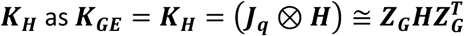 with: ***K***_***G***_ = ***J***_***q***_ ⊗ ***G*** and ***K***_***E***_ = (***Φ*** ⊗ ***J***_***p***_). We considered that there is no need to replace ***G*** with ***H*** in those effects, as ***H*** is a matrix dedicated to model only the G× E. Thus, this is a solution for incorporating genotype-specific sensibilities into a more parsimonious kernel-based model. The single-step procedures were done using the *AGHmatrix* package (Amadeu et al., 2016).

### M06: Reaction-norm GBLUP with single-step G×E kernel from EPA

Model M06 involves the implementation of single-step weighted G×E kernel considering all the information from the matrices ***G, R, γ*** and ***Φ*** previously described. The goal of this approach is to implement an “environmental learning” practice capable of merging the past trends of reaction-norm (***R, γ***) with expectations of reaction-norm (***G, Φ***). In this model, the main genetic effects and environmental variations are modeled the same way as in models M04 and M05, that is, as: ***K***_***G***_ = ***J***_***q***_ ⊗ ***G*** and ***K***_***E***_ = (***Φ*** ⊗ ***J***_***p***_), respectively. The difference in this model is how the nonlinear G×E interaction terms are modelled. This approach is implemented in three main steps.

#### Step 1: Estimating past kernel-realized reaction-norms

First, we estimate the past G×E, that is, the G×E observed in the historical field trials used as training sets. We consider a realization of these past or historical G×E by combining the GRM based on the ***R***-matrix (*t* x *t*) and an ERM based on the ***W***-matrix (*j* x *j*), in which *j* is the number of environments in the training set. Obviously, this step is preceded by the PLS approach, as done for model M05, in which we already estimated the ***R***-matrix, but we also consider a nonlinear **W**-matrix of covariables (***γ***), as in model M03. Using the Kronecker product between these two matrices, we achieve the past G×E (for the historical data set) by ***M*** = ***γ*** ⊗ ***R***, with ***M*** (*s* × *s*) and *s* as the size of historical data using (*s* = *vt*) as the training population set. We used the notation ***M*** to simplify further algebraic demonstrations.

#### Step 2: Estimating future kernel-realized reaction-norms

In an analogy to the method present in the previous subsection, we created a “full rank G×E”, involving all genotypes and locations, as observed in the G×E kernel in model M04. Then we used the static-effect matrix, built up from the environmental weights that each location historically faced (***Φ***) as a precursor of a next-year growing condition, plus a G×E kernel (here referred to as ***N*** to facilitate the mathematical demonstration) created as: ***N*** = ***Φ*** ⊗ ***G***, with ***N*** (*n* × *n*) as *n* is the number of genotypes and environments considered for analysis (training set + testing set). We used the notation ***N*** to simplify further algebraic demonstrations.

#### Step 3: Combining past and future kernel realizations of G×E

Finally, the end-result weighted kernel for G×E effects (***K***_***GE***_) is directly computed by combining the ***M*** and ***N*** kernels as:

If ***M*** = ***γ*** ⊗ ***R*** and ***N*** = ***Φ*** ⊗ ***G***, then:

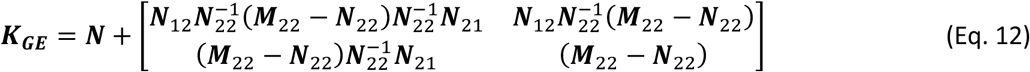

with

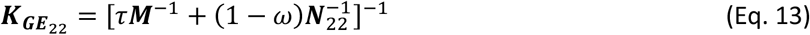

where ***K***_***GE***_ is an *n* x *n* dimension, and *τ* = *ω* =1. In Eq. 12, we have two groups of genotype-environment combinations. Group 1 involves the genotypes already evaluated in past field trials, for which we have all possible information related to the phenotypes, reaction-norm and faced environmental growing conditions. Group 2 is based on genotypes never tested in any past field trial, and environments for which we only know the static-effect priors obtained from long-term historical envirotyping analyses. All analyses were conducted using R statistical software (R Core Team, 2022).

### Prediction scenarios over the historical CIMMYT wheat trials

Four scenarios were evaluated in this study. **Scenario 1**: the first scenario considered the predictions of an entire new year using the previous year as a training set. As the data set is highly unbalanced, this also involved the prediction of new genotypes at locations that may or may not be considered in the training set. **Scenario 2** considers the same approach, but a different model was adjusted for each one of the three TPE previously identified by Crespo-Herrera et al. (2021) (see the Historical wheat trial data section). **Scenario 3** is the same as Scenario 1 but considers multiple years as the training set. In this scenario, we considered two consecutive years to predict a subsequent year. **Scenario 4** is the same as Scenario 2, but also considers pairs-of-previous years as a training set for a subsequent year, but for each TPE individually. The goal of Scenarios 3 and 4 was to verify if it is necessary to use phenotypic data from more than one subsequent year because subsequent years are expected to have the highest genetic correlation, in addition to being more similar in environmental conditions. Thus, the comparison of Scenarios 1-2 and 3-4 may support the interpretation of how phenotypic and envirotypic information from another TPE is useful for making broad predictions for a following year, or whether the use of TPE-specific models is more suitable.

Three statistics were used to measure the quality of the prediction: (1) average predictive ability (*pa*) at “model level”, as is conventionally done for testing models in genomic prediction; (2) genotype-specific predictability, based on the Spearman rank correlation between the predicted and observed values for each genotype in a new year, as suggested by each genotype (Costa-Neto et al., 2021a,b); (3) the relation between the predictive ability of the model and the proportion of the wheat population that was considered “predictable”, that is, with *pa* ≥ 0.2. Here we consider the combination of predictive ability and resolution as a practical measure of the accuracy and usefulness for plant breeders when making decisions for the following years.

### Search for Essential Locations in Indian TPEs

The last proposal of our theory relies on the supposition that a good enviromic-based pipeline for testing cultivars and training models for predicting G×Y must also account for the search of “essential locations for phenotyping”. This approach was implemented by running a grouping analysis of the locations using the environmental weights (***φ***) instead of using raw envirotyping data as was done in the past. For this, we ran a principal component analysis (PCA), followed by a hierarchical clustering using the Euclidean Distance followed by the Ward’s grouping algorithm. These approaches were implemented using the package FactorMineR (Lê et al., 2008). Then, a number of clusters (Nc) found in this analysis was used as a to the number of “essential locations”.

Next, we searched for the most representative locations at each cluster, here called “pivot locations”. Again, the matrix of environmental weights (***φ***) was used for this purpose and were dissected using a genetic algorithm based on PEVmean criteria, as implemented in the SPTGA R package (Akdemir and Sánchez, 2019). The genetic algorithm was parametrized using 100 iterations, and five solutions selected as elite parents were used to generate the next set of solutions and mutations of 80% for each solution generated. For each cluster, we analyzed the similarity between each location and its cluster-pivot location in order to find possible “replacements” for those locations. This approach was done by selecting those locations with a similarity equal to or higher than 95% with each cluster’s pivot location. The clusters found in this study were also interpreted using the TPE characterization done by Crespo-Herrera et al. (2021).

## Data availability

Codes and data are also available at https://github.com/gcostaneto/EPA-PLS.. Codes for creating the matrices and genomic predictions are available at https://github.com/gcostaneto/EPA-PLS. The other codes are available upon request from the first author.

## RESULTS

### Variable importance for modeling environmental similarities across years

This section presents the results of the environment-wide characterization focused on understanding the envirotype-phenotype association (EPA) driving similarity among trialing sites (locations). The results are presented (and interpreted) in two steps: (1) in terms of variable importance in projection (VIP) and (2) the use of weights for grouping environments and finding essential locations for METs in India.

Figure 1 shows the general characterization of the 36 locations in India using the seasonal envirotyping covariables and EPA analysis across years. The relative values of VIP % derived from the multivariate PLS (PLS 2, Figure 1A) involving the long-term envirotyping for locations and an empirical environmental index (from phenotypes). The number of latent vectors (LV) was equal to 8, which was selected based on the number of LV capable of explaining at least 90% of the phenotypic-based environmental similarity. For computing the most important variables, we assumed a threshold of 95% (here relative to VIP % equal to or higher than 85%). For identifying key development stages, we ran the same analysis for each panel (development stage or soil properties) to find the average VIP for each panel (Figure 1A).

**Figure 1.**
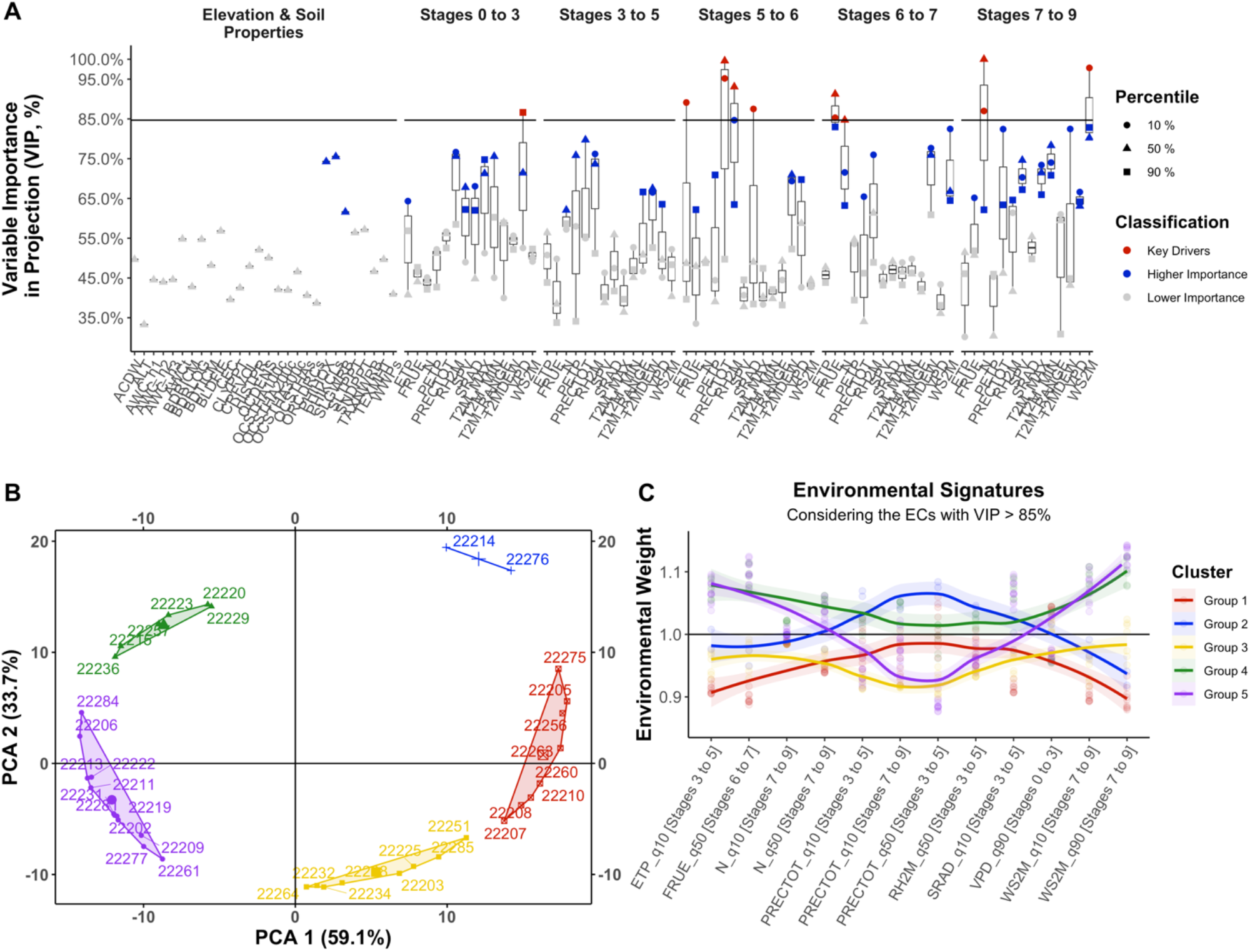
Environmental characterization of the CIMMYT’s Indian locations using EPA analysis covering the seasonal effects across 36 locations over the years between 2011 and 2018. **(A)** Summary of the relative variable importance on projection (VIP) in defining the environmental similarity across locations, considering for it the main environmental variables and development stages (panels) after quality control. A horizontal solid line indicates the threshold of VIP % = 85%, which corresponds to the absolute value relative to the 95% of probability in the distribution of computed VIP values. **(B)** Principal component analysis and clusters found by the Ward Algorithm using the environmental weight as covariables; **(C)** Environmental signature of each group found considering the environmental weights ECs with VIP > 85%. Detail about the all signatures for all ECs are given in Supplementary Figure 2. Environmental weights equal to 1.0 (horizontal solid line) denotes that the given EC is not weighted by EPA. Each dot represents the actual weight of the respective EC for a given a location within the group. The smooth line represents the overall trends considering the key drivers (red dots found in the label A).

From the environmental weights (***φ*** matrix), it was possible to group the location in 5 groups (Figure 1B). These groups were found by using the matrix for a principal component analysis (PCA), followed by a hierarchical grouping using Ward’s algorithm. Then we were able to pinpoint the ECs with higher importance (VIP > 85%) as a descriptor of the environmental profile that each group has (Figure 1C). However, our results suggest that every environmental factor has a minimum VIP% of ∼30% (Figure 1A), which means that not one piece of environmental information is irrelevant, but some of them are responsible for the main expected similarities between the groups of environments (Figure 1C). As a great number of environmental factors have greater importance (60-85%, green colors, Fig. 1A), all ECs were used to identify the group environments.

With regards to development states, the stage from heading to anthesis (stages 3 to 5) is key for differentiating the quality of the growing environments for wheat in India. At this stage, a great number of variables has the highest impact (VIP >=85%), such as the minimum quantile of evapotranspiration (ETP at q10), median value of air humidity (RH2M q50), minimum quantile of global solar radiation (SRAD at q10) and rainfall precipitation (PRECTOT q10). Other important combinations of stages/variables include the vapor pressure deficit (VPD) at the early vegetative stages (e.g., during double ridge appearance), as the impact of temperature on radiation use efficiency (FRUE) at anthesis (stages 6 to 7) are key factors contributing to increasing the diversity of growing conditions.

### Locations that represent the diversity of growing conditions in India

Then, we used the number of clusters (5) as an indication of the effective number of locations that represents the environmental diversity of the locations in CIMMYT’s experimental network for wheat in India. Next, we used a genetic algorithm based on PEV mean criteria to analyze the matrix and identify the five locations most likely to be “pivotal-locations” for the phenotyping network for India. In this analysis, we identified locations ID: Group 1 = 22231, Group 2 = 22257, Group 3 = 22260, Group 4 = 22268 and Group 5 = 22276 (Figure 2). This means that these locations are essential for conducting adequate phenotyping that is capable of capturing the diversity of growing conditions, thus lowering the number of field trials.

**Figure 2.**
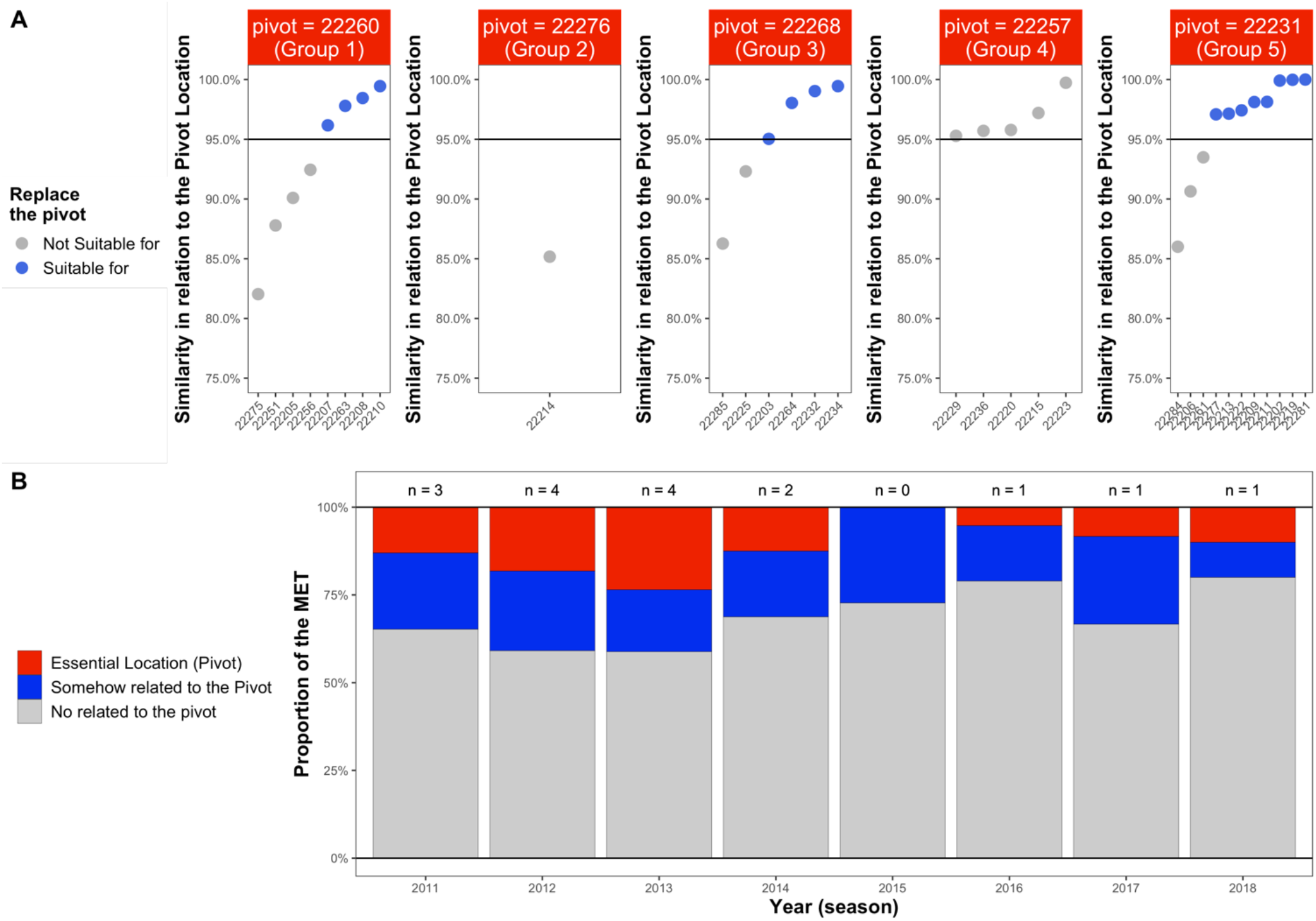
Diagnosis of the environmental diversity observed within each cluster of locations found in this study for CIMMYT’s wheat trials in India. **(A)** Relative similarity between each location and its pivot-locations within each cluster. Each pivot was selected by the clustering analysis of the environmental weights from the EPA analysis using the multivariate PLS 2 algorithm. **(B)** General overview of the representativeness of the multi-environment trial (MET) network in India for each season (year). Values above (n=3, n=4, etc.) indicate the number of pivot locations considered in each year. Red colors denote the proportion of essential locations (pivot) in each year. Blue colors denote the proportion of environments somehow related to some pivot (similarity >= 95% with the pivot), using as threshold 95%. Gray colors denote the proportion of environments considered to represent the diversity of the MET.

As occurs in any real-world breeding program, due to some external factor (logistic, political issues, budget), some favorable locations are not available for conducting future field trials. To avoid any theoretical gaps in the methodological analysis, we proposed a score of the locations in relation to the pivot location of each group identified in the clustering analysis (Figure 2). As a threshold, we adopted a similarity equal to or higher than 95%. We observed that the following locations (gray colors) are not necessary for the experimental network: group 1 (22275, 22251, 22205, 22256), group 2 (22214) and group 5 (22284, 22206, 22261). Location 22276 (group 2) seems to be essential, and no other location is capable of replacing it. For Groups 4 (pivot 22257) and 5 (pivot 22231), it is easy to replace this location with any other, giving the opportunity to use other locations in this group for screening secondary growing conditions, such as fertilizer or irrigation levels and biotic stresses.

Based on the TPE to which each one of those pivot locations belongs, there is a concordance with the study conducted by Crespo-Herrera et al. (2021). However, variations within each TPE were detected, which increased the resolution to find key locations that represent the variability of all MET in India. For example, the diversity of TPE 1 is represented by two pivot locations, 22260 (group 1) and 22231 (group 5), where it seems that the patterns the ECs ETP_q10 [Stages 3 to 5], FRUE q10 [Stages 6 to 7] and windspeed (WS2M_10 and q10 at stage 7 to 9) where key for splitting TPE 1 into two distinct groups (Figure 1C). The diversity of TPE 2 is represented by the pivot location 22257 (group 4). Another within-TPE division was observed for TPE 3, which is now represented by groups 22276 (group 2) and 22268 (group 3). Within TPE 3, the locations that represent the diversity are differentiated by ECs for a specific development stage and with a major effect on the atmospheric water balance, mostly related to precipitation, radiation and air humidity during Stages 3 to 5.

Hence, the design of field trials in India must follow the pivots (and the environments related to these pivots), as described in Figure 2, where it should be possible to better represent the G×E for the purposes of data analysis and training prediction. It is interesting to note that TPE 1 is the easiest to design, as we have two main pivots and a larger number of highly similar locations as options. In TPE 3, the same situations occur, but one essential location (22276) seems to be indispensable to better capture the diversity of TPE 3. It seems that TPE 2 is the easiest to design due to the higher similarity among the locations that are part of it. Thus, the characterization done by Crespo-Herrera et al. (2021) proved to be very successful for gathering the locations within TPE 2; however, there is a remaining diversity in TPEs 1 and 3 that is now better understood due to the current study. The ideal MET must consider the five pivot locations in each year. However, this is not the situation observed in the Indian MET considered in this study (Figure 2B). The implication of this condition is discussed in the diagnosis of the joint predictive ability trends across years in the next sections.

### Plasticity patterns reveled by the univariate PLS 1 algorithm

The reaction-norms were computed for each genotype using the univariate version of the PLS algorithm (PLS 1). To run this algorithm, we first assumed the ideal number of latent vectors (LV) as those capable of explaining at least 90% of the genotype-specific G+GE variability. After a basic study and a data control analysis, we identified 61

ECs suitable for this analysis (Figure 3), which means that ECs with missing values in some environment or with difficult to interpret biological meaning (e.g., reaction-norm for wind speed) were removed in this step.

**Figure 3.**
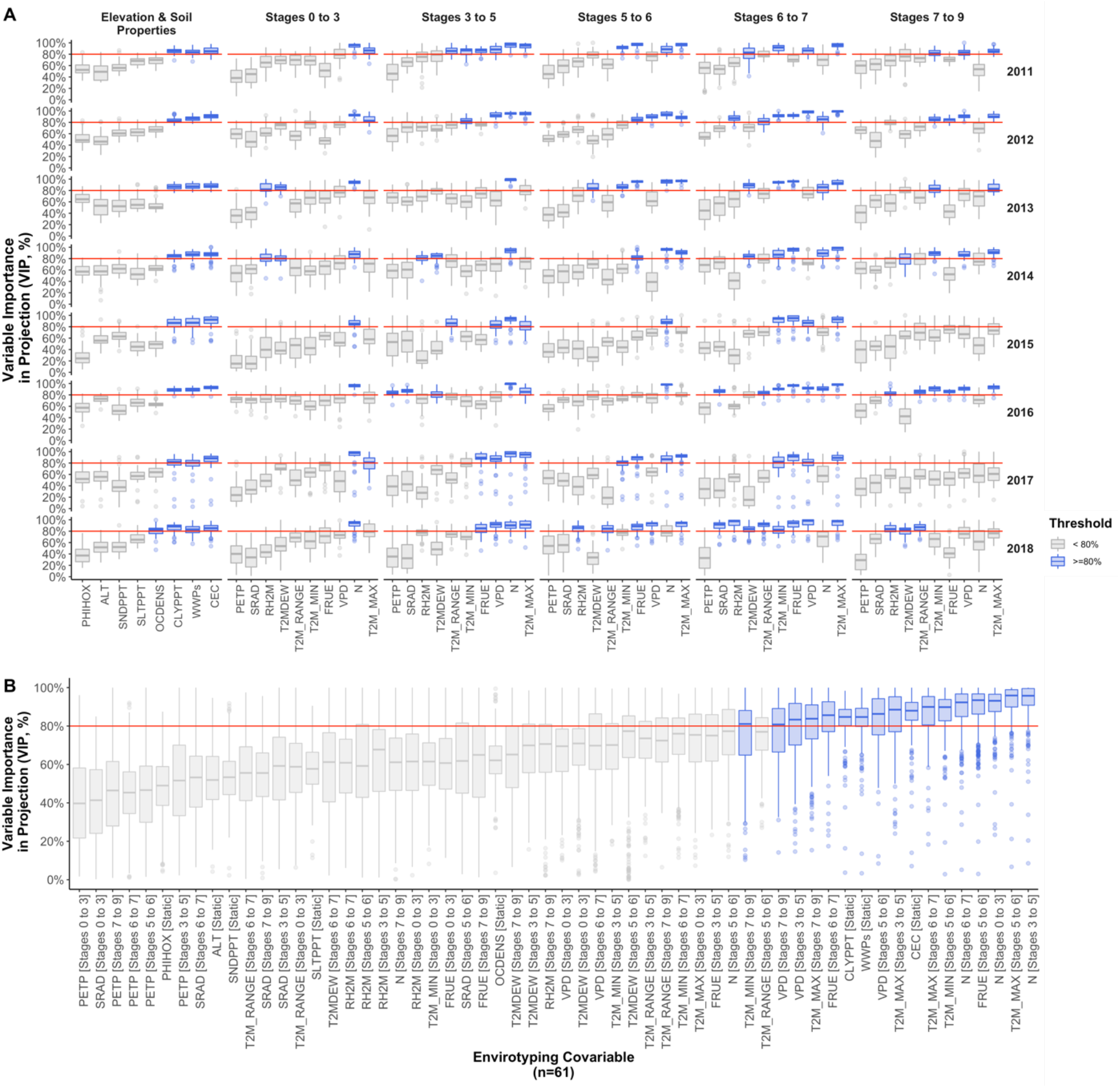
Distribution of the relative variable importance in projection (VIP, %) from the PLS 1 analysis for each genotype considering each envirotyping covariable (environmental factor x development stage) and years between 2011 and 2018. (**A**) detail of VIP distribution considering all genotypes and detailing for each year and development stage. (**B**) general overview considering all years. The vertical red line denotes the VIP = 80%. Blue denotes those variables with higher importance (VIP > 80%) for the given year/development stage combination.

A major proportion of the genotypes were well fitted (R^2^ ≥ 0.90) with the number of latent vectors ranging from 3 to 10 (>90% of the germplasm), and a few were outliers, such as LV from 15 to 30 (2% of the germplasm).

Then we collected the variable importance in projection (VIP) statistic and converted into relative VIP % by dividing the VIPs for each genotype by its maximum VIP value (that is, the most important environmental variable for a given genotype). The distribution of the % VIPs considering all genotypes is given in Figure 3.

Soil properties such as CEC (soil cation exchange capacity), WWP (soil water wilting point) and CLYPPT (clay content) seem to be essential for modeling the adaptability of the genotypes in all years considered (Figure 3A). For the meteorological variables (divided by development stages), we observed a wide range of VIPs (sometimes ranging from 0% to 100%), which indicates huge variability in the germplasm in terms of reaction-norm. Taking the average VIP for each variable and looking at those higher than 80% (that is, of higher importance), this variability is also reflected in differences among years and crop development stages. Some variables exhibit a consistent higher VIP% across years for most of the germplasm (Figure 3B), such as the photoperiod (N) in the early vegetative stages, such as 0 to 3 and 3 to 5. Factors related to the air temperature such as maximum temperature (T2M_MAX) in stages 5 to 6, 6 to 7 and 7 to 9, in degree of importance, respectively, seem to play an important role in the plasticity of the wheat germplasm in India. The temperature range (T2M_RANGE) is also important for stages 7 to 9. Finally, a related covariable is the effect of temperature on radiation use efficiency (FRUE), which is highly important for reproductive stages 5 to 6 and 6 to 7, but has a minor effect during the vegetative stages. Another important EC is the variables of vapor pressure deficit (VPD) during reproductive stages (5 to 6), during terminal spikelet formation (3 to 5) and grain filling to maturity (7 to 8).

The diversity of genotype-specific coefficients (***Λ*** matrix, illustrated in Figure 4A) highlights the problem of selecting environmental covariables that explain G×E. There is huge variability in signal (positive/negative reaction-norms) and magnitude across different years and germplasm (as each year has a different set of genotypes). A few genotypes demonstrate “outliers” in their reaction-norm responses for some specific factors (extreme blue or red colors). On the other hand, most of the reaction-norms range from -0.5 to 0.5 (orange to green), but their diversity across diverse development stages and years suggests the difficulty of selecting a few covariables to really explain the G×E across years. Because of this, we considered the matrix of all genotype-specific coefficients (***Λ***) to create the **R**-matrix (Figure 4B). In this **R**-matrix, it seems that there is a second genetic relationship matrix that takes the shared reaction-norm patterns observed in past field trials as a sign of the expected genetic-based future joint G× E variations (Figure 4B).

**Figure 4.**
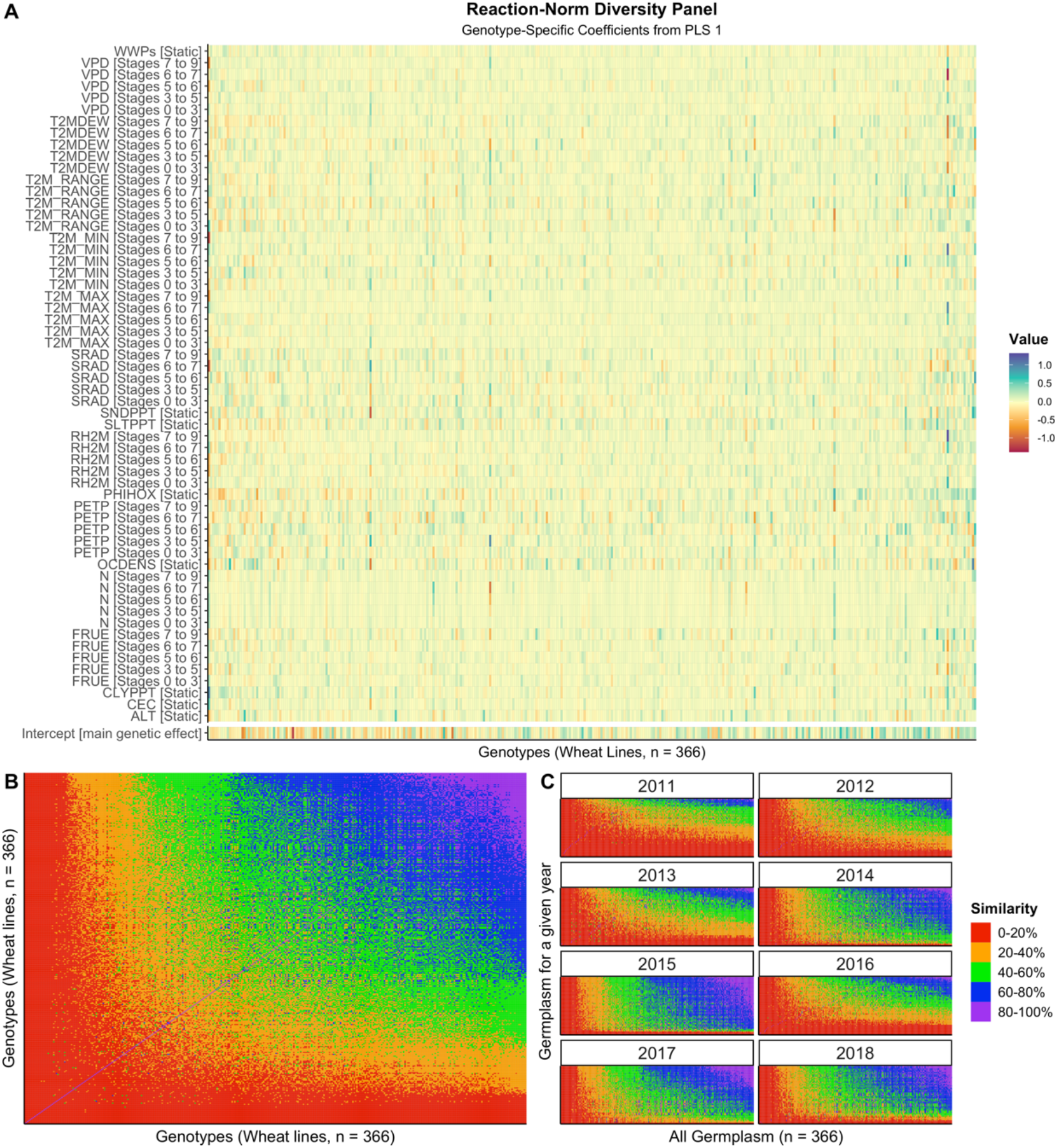
Characterization of the reaction-norms the subsequent similarity of the plasticity patterns for CIMMYT’s wheat germplasm in India. **(A)** panel of genotype-specific coefficients for each genotype (columns) and envirotyping covariables (rows) across from the EPA analysis over the G+GE effects across years; **(B)** genetic similarity (gaussian kernel) of the empirical plasticity computed from the reaction-norm coefficients treated as “plasticity markers”. **(C)** patterns of similarity between the germplasm tested in a given year (from 2011 to 2018) and the all germplasm (n=360).

This empirical relationship can also be useful to understand the lack of relatedness between the set of genotypes (wheat lines) in a given year and the whole breeding germplasm– in this case, the Indian wheat germplasm from CIMMYT (Figure 4C). The set of genotypes evaluated in 2015, for instance, is highly related (blue-purple colors) to the core germplasm, which can suggest, for example, that this germplasm is a good training set for predictions across years. On the other hand, the set of genotypes in 2011 is poorly related to the core germplasm (black, red colors), which is also a sign of the lack of genetic relatedness and the differential genetic progress observed in the Indian TPEs that have shaped (and diverged) the past wheat lines from the more recently developed germplasm. For prediction purposes, we considered the **R**-matrix for pairs of years (or three years, according to the prediction scenario) for running our genomic prediction analysis.

### Comparison between linear and nonlinear environmental relationships for G×Y predictions

The average values of predictive ability, measured by the Spearman correlation, are illustrated for each model and scenario in Figures 5 and 6. As expected, the baseline model without enviromics (M01) was the worst model for any scenario. The average predictive ability for this model (horizontal red lines) was very close to 0 (horizontal black line), below 0 for the analysis considering all TPEs together (Scenarios 1 and 3) and above zero for those models fitted for each TPE independently (Scenarios 2 and 4). This result suggests that the definition of TPEs, followed by fitting independent models for each TPE, is a good strategy when no enviromic data are considered. TPE 2 was the most predictable without enviromic data, but even in the best scenario, a lower predictive ability (r ≅ 0.20) was observed, and in most cases, r ≅ 0.00 or negative. In Figure 6B, we present a fair comparison between how frequently the alternative models outperform M01.

**Figure 5.**
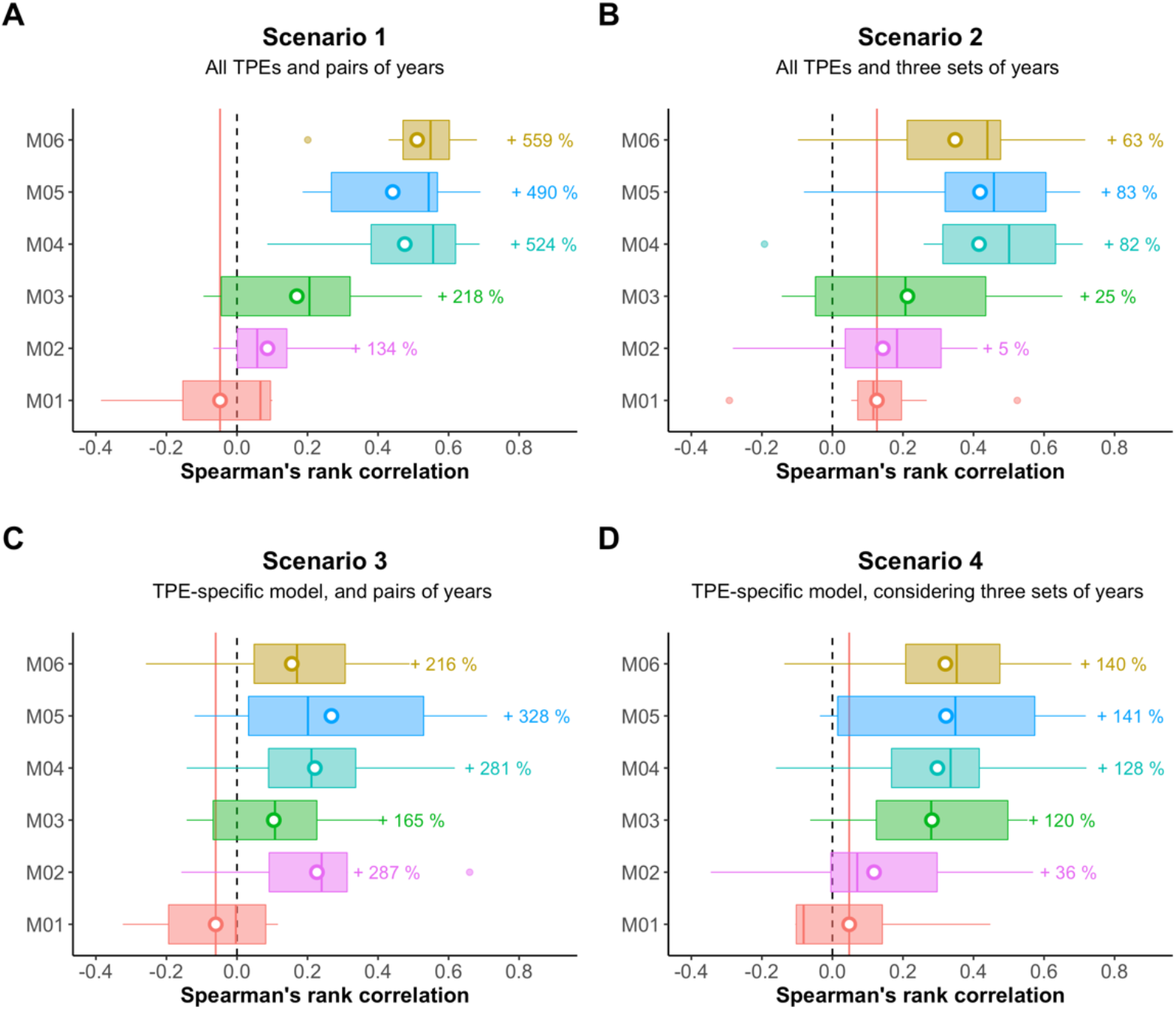
Predictive ability for grain yield considering each statistical model and prediction scenario of G× Y between 2011 and 2018 in India. (**A**) <u>Scenario 1</u>, considering all TPEs together and a pair of years (the previous year as the training set, the following year as the testing set). (**B**) <u>Scenario 2</u>, considering all TPEs together and three sets of years (pairs of subsequent previous years as the training set, a single following year as the testing set). (**C**) <u>Scenario 3</u>, for each TPE, considering a pair of years (the previous year as the training set, the following year as the testing set). (**D**) <u>Scenario 4</u>, for each TPE, considering three sets of years (pairs of subsequent previous years as the training set, a single following year as the testing set). The white points within each boxplot represent the average value (mean), while the horizontal lines represent the median (quantile 50%). Vertical lines denote the reference of not predictability (null predictability = 0, dashed black line) and the reference of the average predictability in terms of the Spearman rank correlation of the baseline model (M01, no enviromics, red line). Percentage values above each boxplot represent the average gain/loss in predictive ability in relation to the baseline M01

**Figure 6.**
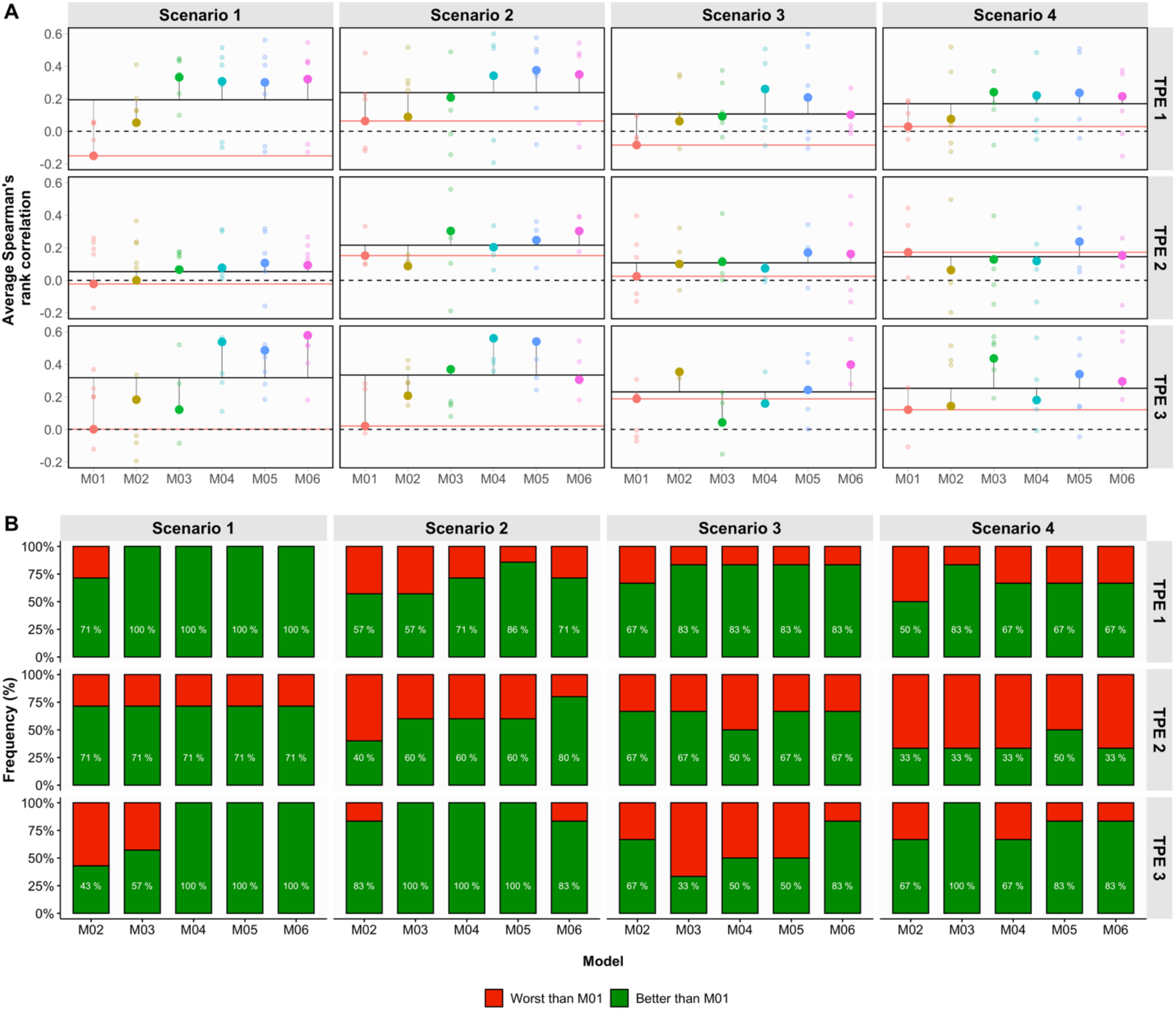
Summary of the TPE-specific predictions considering the years between 2011 and 2018 in India. (**A**) Average predictive ability (PA) for grain yield for each statistical model and prediction scenario. Vertical lines denote the reference of not predictability (null predictability = 0, dashed black line), average predictive value for each panel and TPE, considering all models (solid black lines), and the reference of the average predictability of baseline model (M01, red line). (**B**) head-to-head relative frequency of the PA superiority of each model (M02-M06) over the baseline GBLUP (M01) across years, with “better than M01” (green colors) indicating how many times each one of the tested models (M02-M06) outperformed M01 and “worst than M01” (red color) relating on how many times the M01 was the winner model.

In this study, our goal was to predict a new year, but before going any further, it is essential to discuss models M02 (conventional linear reaction-norm, Jarquín et al., 2014) and M03 (nonlinear reaction-norm, Costa-Neto et al., 2020b). On average, these models outperformed the M01 models in every scenario (Figure 6A), with gains up to +218% (Scenario 1, M03) and +287% (Scenario 3, M02). For any TPE and scenario, the M02 and M03 seem to always outperform the M01 in most years, except for Scenario 1 – TPE 3 (M02) and Scenario 3 – TPE 3 where the use of M01 fitted using TPE-specific environments seems to lead to better results (Figure 6B). Both models are based on the same W-matrix of covariables (the same used to compute the genotype-specific coefficients in the PLS 1 step). However, the difference is that one model accounted for a linear approach for the ERM (correlation based) and the second model considered the nonlinear approach (Gaussian process for the similarities). Except for Scenario 3 (Figure 5), in all other scenarios the M03 outperformed M02, both in terms of higher mean (dots) and median (horizontal colorful lines). Figure 6 indicates that in Scenario 1 (using two years of data in a single model for all TPEs), the gains compared to M01 observed for M03 ranged from +90% (TPE 2) to +480% (TPE 1), while for model M02, they ranged from +20% (TPE 2) to +200% (TPE 3). A similar trend was observed for Scenario 2, in which in TPE 2, the M02 was outperformed by M01 (−15% reduction), while for the same TPE, the M03 outperformed M01 by +37%. For Scenarios 3 and 4 (using more than 2 years), the same trend was observed for those three models, except for TPE 3 in Scenario 3, where M03 was the worst model. Thus, in general, the use of a nonlinear kernel for ERM (M03) outperformed the conventional way of using environmental information in genomic prediction (M02).

### Accuracy gains when adding EPA as an intermediary step in GBLUP

Looking at only the benchmark models (M01, M02 and M03), the results suggest that TPE 1 was the TPE that benefitted the most by the inclusion of enviromics, irrespective of the kernel method or envirotyping protocol used. However, when we analyzed the proposed enviromic-aided strategies (M04, M05 and M06), the gains were outstanding for most scenarios and TPEs. For example, in Scenario 1 (Figure 5), while M03 achieved a gain equal to +218% (r =0.18) over M01 (r=-0.05), the replacement of the conventional ***W***-matrix by the environmental weights from the S-matrix (M04) achieved a gain of up to +524% (r=0.58). Higher gains from M04 were also observed in Scenario 2 (+82%, r = 0.41), Scenario 3 (+281%, r=0.25) and Scenario 4 (+128%, r = 0.33). Considering the predictions for each TPE (Figure 6A), model M04 was very effective for predicting the TPE 3 in most scenarios, with gains due to M01 up to +540% (Scenario 1, r =0.54), +386% (Scenario 2, r=0.56) and +18% (Scenario 4, r = 0.18); however, there was a reduction of -6% for Scenario 3 (r=0.16). A negative result was also observed for TPE 2 in Scenario 2 (+12%, lower than other models such as M03) and Scenario 3 (−11%). Thus, model M04 aggregates benefits and increases accuracy over M03 in most cases, but for some scenarios/TPE, we observed instability in the predictions due to a lack of correlation relative to the “genetic causes of G× E”, that is, the environmental weights alone generally help to increase the prediction, but they are not enough to ensure stability in the predictions across all scenarios and TPEs.

The inclusion of the reaction-norm based matrix (**R**-matrix, M05) helped solve these issues in some scenarios and TPEs (Figures 5-6), but it does not always add accuracy in relation to the standard use of the environmental weights. For example, the comparison of the observed global predictive ability (Figure 5) for models M04 and M05 shows almost the same gains for the scenarios that include two-year data (Scenarios 1 and 2), but higher gains for scenarios that include more than one year (Scenarios 3 and 4). In this last scenario, the gains over the M01 ranged from +328% (r = 0.28) and +141% (r=0.27), respectively. The predictions for each TPE (Figure 6A) were also interesting, achieving the best performance in terms of predictive ability in Scenario 2 for TPE 1 (r= 0.38, +145% over M01) and TPE 3 (r=54, +371% over M01). Another interesting comparison is observed in TPEs 2 and 3 in Scenario 3, with gains ranging from +10% to +88% compared to -6% to 25% observed for M04. For TPE 1, however, it seems that M04 and M05 are always better choices for modeling in any scenario in comparison with M01 (Figure 6B), while M06 seems to present almost the same performance despite being a more complex modeling approach.

Finally, model M06 is the single-step combination of the reaction-norm matrix and environmental weights. In most cases, the models seem to achieve similar performance as their predecessors (M04 and M05), but when looking at the variability of the predictive abilities (size of the boxplots, that is, 25% to 75% quantiles), these models suggest more consistent predictions across all scenarios (Figure 6B). Model M06 achieved its highest performance in Scenario 1, with gains ranging from +130% (TPE 2) to +460% (TPE 1) and +540% (TPE 3). For TPE 3 in Scenario 3 (Fig. 6A), this model was the best model (r=0.40, +44% over M01), with potential to achieve predictive ability values close to r = 0.65.

### Resolution of the G× E prediction at the genotype level

The resolution of the G×E prediction was given by the genotype-specific Spearman correlation between observed and predicted yield values for a future year. Figure 7 presents these results considering all years (from 2012 to 2018; 2011 is not considered because it was not predicted). For all models, the differences in the prediction scenarios affected the resolution of the G× E prediction. The resolution of M01 (no enviromics) ranged from 3% (Scenario 4) to 35% (Scenario 2). For predicting a new year based on the previous one, this resolution was equal to 11%. This value means that the genomic prediction model is capable of adequately predicting the G× E variations for 11% of the germplasm (the remaining 89% is unpredictable). The inclusion of enviromics was key for increasing the resolution. For Scenario 1, the use of a nonlinear kernel for **W**-matrix (M03) pushed the resolution from 11% (M01) and 25% (M02) to almost 50% of the germplasm. This same trend was observed for Scenario 4. However, when a TPE-specific model is fitted (Scenario 2), those three benchmark models had almost the same proportion of the germplasm predicted (∼45%). It should be noted that the resolution of the M03 was still the highest one (proportion of blue and purple colors). In Scenario 3, M02 exhibited the highest resolution, perhaps due to the fact that in this case the linear kernel adequately captured the phenotype-envirotype association driving the G× E pattern population set.

**Figure 7.**
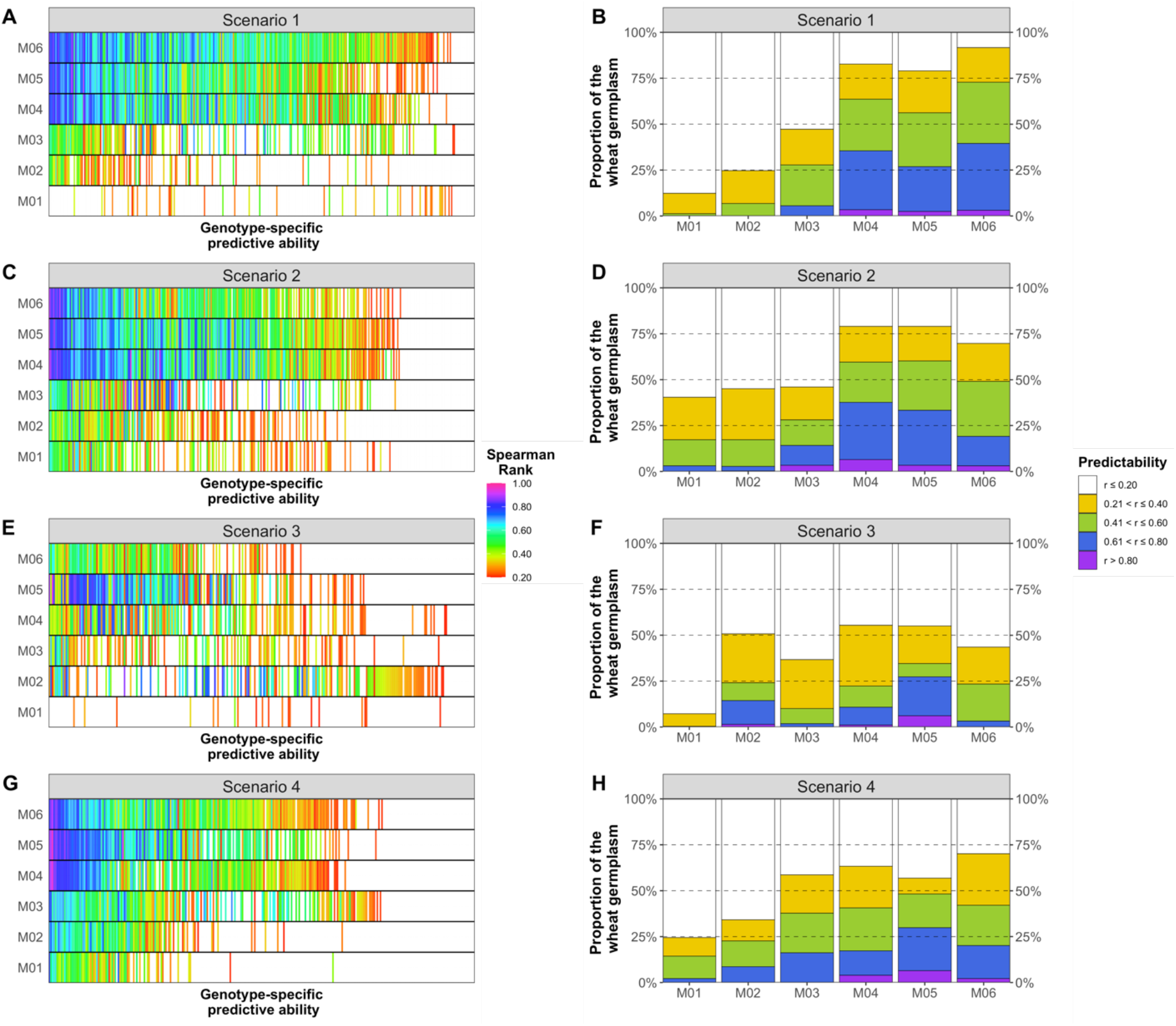
Resolution of the genomic prediction models for each scenario tested. (**A**) heatmap of the spearman’s rank correlation for each genotype in prediction Scenario 1. (**B**) Resolution in terms of the distribution frequency of predictability for Scenario 1. (**C**) Heatmap of Spearman’s rank correlation for each genotype in prediction Scenario 2. (**D**) Resolution in terms of the distribution frequency of predictability for Scenario 2. Heatmap of Spearman’s rank correlation for each genotype in prediction Scenario 3. (**F**) Resolution in terms of the distribution frequency of predictability for Scenario 3. (**G**) Heatmap of Spearman’s rank correlation for each genotype in prediction Scenario 4. (**H**) Resolution in terms of the distribution frequency of predictability for Scenario 4.

The highest resolution was achieved by model M06 in prediction Scenario 1. This means that in this scenario, we observed the highest predictive ability gain (+559%, r = 0.56, with r_max_=0.69) and highest resolution (∼95% of the germplasm). This resolution was drastically reduced in the other scenarios, especially 2 and 3 (inclusion of more than one year), which suggests that the increase of phenotypic data does not act as a source for increasing the resolution of G× E. The separation of the models for each TPE using two-year data (Scenario 2) seems to be a satisfactory strategy for most models, but here models M04 and M05 outperformed model M06. This suggests that a good adaptability pedigree within each TPE and also a good measure of the within-TPE weights must be useful for training the models.

## DISCUSSION

In this study, we present results addressing the following topics: (1) how nonlinear kernel methods seem to outperform linear methods in terms of accuracy; (2) how the enviromics linking EPA studies could leverage the current TPE characterization priors used to fit TPE specific GBLUP models, while borrowing information within and across TPEs; (3) how environment-wide characterizations revealed the differential effect of the seasonal variations in magnitude, direction and impact of the environmental factors in the trialing networks across different years, which affects any reaction-norm estimation; (4) how EPA studies could help the development of reinforcement algorithms for “learning” how the macro-edaphoclimatic variations affect the environmental quality for a given trait across historical data sets and ; (5) how the current cultivar testing pipelines can accommodate these EPA approaches into a breeder’s friendly manner, without adding costs to the current budgets, and how this could add value in the decision-making processes of selection and design of more representative and cheaper trialing networks.

### EPA enables environmental characterization and plasticity modeling

In this study we introduce a pipeline for characterizing the similarity among locations using EPA, envirotyping data and historical yield data. In summary, our environment-wide characterization suggests that: (1) soil properties and elevation are less important for describing the variation across years (average VIP % = 50%, with maximum average higher importance equal to 60%), which is reasonable when considering that the main difference observed in environmental conditions is due to seasonal variations in meteorological variables; (2) the temporal variation of meteorological factors is key for understanding the similarity among environments, in which some variables are more important than others due to the crop-specific developmental stages; and (3) due to the use of diverse statistics for capturing the distribution of these factors, we observed that median values are not always the best statistics for describing the distribution and the weight of some environmental factors. Hence, we envisage that the process of selecting covariables is not the ideal situation for modeling the effects of seasonality across years, because the envirotype-phenotype patterns are not static across years due to seasonal variations in climatic factors and genetic variations between the different genotypes evaluated in each cycle. Thus, although the EPA approach was useful in revealing some “environmental signatures” that describe the major factors affecting the diversity of growing conditions within TPEs, the use of environmental-wide factors seems to be a more feasible and conservative way of modeling the similarities among locations in the future.

The EPA analysis of genotypes (reaction-norm modeling) also suggests that the selection of covariables as part of prediction models could be a contradictory strategy. While the selection of covariables could increase the accuracy of previous years, which can be useful for exploratory purposes, it may not be feasible for supporting prediction models in real-world breeding programs, especially when the goal of these models is the prediction of future years (new G×Y) where the plasticity of the yet-to-be-seen genotypes is unknown. This is a consequence of the huge time-scale differences in VIP% across years (seasons), germplasm (different genotypes evaluated in different seasons) and across development stages. Because of this, for further analysis, we advocate the using all envirotyping covariables for creating the environmental relationship matrices – and consequently, the kernel-based reaction norms in genomic prediction. For modeling reaction-norms, here the PLS approach seems to simplify the diversity of reaction-norms too much; consequently, we suggest that in future studies, nonlinear approaches such as GAM (Heinemann et al., 2022) be used as an alternative approach. Despite this, the use of PLS-based genotype-specific coefficients of reaction-norm were successful in recovering the main trends of plasticity and leverage of the genomic prediction for future years in the historical data sets. This will be discussed in further sections and has been observed in other applications described by Montesinos-Lopez et al. (2022).

### Linear kernel methods do not capture the nonlinearity among field growing conditions

The statistical modeling approaches for the analyses of plant breeding in multi-environment trials continue evolving as more data with more complex structure are being collected in plant breeding programs (Crossa et al., 2021, Teixeira et al., 2011). For example, diverse set of methods are available for dealing with multi-dimensional data, such as the nonlinear approaches that are now being applied for modeling environmental relatedness using large-scale envirotyping data (Washburn et al., 2021; Rogers et al., 2021; Westhues et al., 2021; Costa-Neto et al., 2021a,b). Here were compared the conventional multi-environment GBLUP (M01, no enviromics) and two reaction-norm GBLUPs, the first using envirotyping data on a conventional linear kernel (M02, linear W-matrix, Jarquin et al., 2014) and the second using these data on a nonlinear Gaussian kernel (M03, nonlinear W-matrix, Costa-Neto et al., 2020b). M02 and M03 assumed that *we* k*now the future envirotyping data*, but model M03 (nonlinear kernel) outperformed model M03 (linear kernel) in every scenario tested. This result agrees with what was achieved in wheat in Australia (He et al., 2019) and tropical maize in Brazil (Costa-Neto et al., 2020b), in which both the use of nonlinear kernels (Gaussian kernel and/or Deep Kernel) were superior to the linear kernel for modeling environmental relationships from ECs.

From the biological point of view, it seems reasonable to affirm that environmental covariables are non-additive between each other (Costa-Neto et al., 2021c). Also, environmental covariables have higher collinearity and a lack of orthogonality (Heinemann et al., 2022). Because of this, some studies have added the step of “variable selection” (e.g., Millet et al., 2019; Westhues et al., 2021; Mu et al., 2020), which could help in to overcome this issue but could cost a loss of information when trying to predict a yet-to-be-seen GxE, as will be discussed in further sections of this paper. Here we approached these issues by building non-parametric, nonlinear environmental kernels, that take into account (leveraged) some phenotypic data as prior information to adjust bandwidth factors (Costa-Neto et al., 2021b) and that are able to learn hidden nonlinearities underlying the variations between the observed macro-environmental influence (from envirotyping) and the actual phenotypic variation (the resulted GxE). This “environmental learning” approach is a supervised method that goes in the same direction of other approaches used for this purpose, such as convolutional neural networks (Washburn et al., 2021) and gradient boosting machine (Westhues et al., 2021), but with the benefit of dimensionality reduction as the historical GxE and reaction-norm patterns are summarized in two simple matrices (***γ*** and ***Φ***).

Also, we observed that nonlinear kernels are better at handling outliers in the environmental covariables, and are also more conservative in building similarity relations (not covariances). For instance, the Gaussian Kernel for environmental similarity led to a diagonal matrix (diag(GK) = 1), with off-diagonal effects equal to the degree of similarity (from 0 to 1) among environments. Considering the lack of similarity (identity matrix) is the same as assuming that every envirotyping data does not add value in weighting those similarities. In this sense, a good envirotyping protocol must be designed and conducted in order to avoid misspecification of those similarities, while differentiating the effect of different time-scales (e.g., time-windows during crop lifetime, phenology) according to the crop phenology. For example, there is no meaning in using monthly temperature values to model the growing conditions faced by an annual crop such as maize, due to the fact that during 30-day intervals, the crop goes through a wide number of development stages, and for each one of them, the crop will not face the same environment due to the differential needs and sensibility to inputs and stress factors, as evidenced by Rogers and Holland (2022). On the other hand, the use of high-resolution data, such as an hour-scale, may not aggregate value and moreover, may even be a source of noise that will limit the accuracy of the reaction-norm models (Jarquin et al., 2020).

However, our results show that even when we maintain the same envirotyping protocol, the kernel structure choice is key for the successful prediction of G× E. In fact, the lower accuracy results obtained from linear envirotyping-based ERM in this study, and in other publications (e.g., Millet et al., 2019; Rogers et al., 2021; Rogers and Holland, 2022), can be attributed to the fact that they fail to reproduce the true quality of the environment by overestimating or underestimating its quality. For this reason, data analysts should dedicate efforts towards developing a diverse set of “environmental markers” that can be globally used for a certain species or that are germplasm-specific. These markers must be designed and tested to account for actual plant ecophysiology interactions, which can be computed by crop growth models or using frequencies of occurrence of environmental typologies (Costa-Neto et al., 2021c), which in both ways can be leveraged by using environmental-phenotype association when historical data-bases are available for it, especially when we want to predict future growing conditions based on past priors.

### TPE-specific models do not ensure higher accuracy in G× Y predictions

TPE characterization is a key for reconciling breeding goals and expectations with the actual structure available to support the screening for genotypes as candidate cultivars. This also provides information to check the representativeness of the current experimental network – and, sometimes, highlights key aspects to optimize it. One of these key aspects is implied directly in the design of strategies for crossing and selecting more adapted genotypes for key growing conditions, which leverages the genetic gains and ensures the efficiency of the breeding program, allowing breeders to focus on other breeding goals – that is, secondary traits, such as nutritional quality. Thus, it is a sustainable way to measure the quality of the germplasm for particular growing conditions, and consequently, to better deal with future extreme events, such as climate-change panels around the world. Thus, the envirotyping protocols are now integrated into breeding and post-breeding pipelines as a “thermometer of the germplasm adaptation”, by showing the potential phenotypic landscapes (Messina et al., 2018; Costa-Neto et al., 2020a) and for providing evidence about how the environment could affect past genetic gains (Heinemann et al., 2019; Cooper et al., 2021) and the actions that must be taken to achieve the breeding goals in upcoming years (Elli et al., 2020). In this context, envirotyping data, historical data or simulated data from crop growth models have been used to clearly define the limits of each TPE in breeding programs and, as a result, this prior information can be used to control (or exploit) the G× E patterns within homogeneous growing conditions (Windhausen et al., 2012; Crespo-Herrera et al., 2021).

The G× E interaction is a key property of the experimental network, and its magnitude and nature depend on the diversity of the experimental network (Costa-Neto et al., 2021c;). This was successful in reducing the crossover G× E interaction when targeting cultivars and borrowing information across countries for training genomic prediction (Crespo-Herrera et al., 2021). However, for the prediction of new G× Y, our results suggest that models without the division of TPEs favor the EPA study, as the amount of screened growing conditions for each genotype ensures better estimation of the reaction-norms. In addition, the increased number of observed environments also provides a better estimation of the environmental weights, which favors the enviromic-aided models. The use of a particular model for each TPE favored the classic conventional multi-environment GBLUP (M01) (as expected, models M02 and M04 in most cases). However, the accuracy gains achieved by the models accounting for enviromic-aided pedigrees were much higher, for it was observed that accuracy gains were nearly 1,000%; there were also higher resolution gains.

Due to the fact that our data set is highly unbalanced across years, we failed in testing to see whether replication of certain genotypes across years, for a specific TPE, can be useful to optimize cross-TPE predictions over different years. Thus, it is possible that using TPE-specific phenotypes, while accounting for replications of genotypes across years, could be a good balance between the optimization of independent phenotyping efforts within each TPE and an accurate Envirotype-to-Phenotype Association (EPA) analysis, which is the basis of the enviromics-aided pedigrees. In addition, as observed in other studies in maize (Costa-Neto et al., 2021b), it is possible that not every phenotype data point is useful to aggregate information in genomic prediction – in fact, sometimes this is one source of noise that implicates predictive ability losses. Then, the use of multi-year data, under unbalanced conditions, might not contribute at all.

For some years, the lack of similarity present in the ***R*** matrix also partially reflects the lack of accuracy. In the years 2012 and 2016, even when considering more phenotyping data (across years and TPEs), the observed predictive ability was low for most of the models.

### Association studies between envirotyping data and phenotypic variation is key for dealing with the uncertainty of future G× Y

The use of historical yield data accomplished with long-term envirotyping is the basis for understanding how, in reality, the environmental factors (meteorological, soil, biotic factors), across the plants’ lifetime, affects the end-result phenotype of interest. As such, the field of the Envirotype-Phenotype Association (EPA) can approach this uncertainty, by reverting to past studies on factorial regression (e.g., Hardwick and Wood, 1972; Denis, 1988; Costa-Neto et al., 2020a) and partial least squares (Vargas et al., 1999; Porker et al., 2020) to provide a more realistic descriptor of the environmental impacts on the experimental network. As each experimental network is composed of genotypes and environments, two types of analysis were proposed in this study: (1) the search for descriptor adaptability by estimating the empirical reaction-norms of the genotypes tested in the past; (2) measuring the weights of each environmental factor in driving the relatedness and diversity among field trials (or locations across years).

The first approach focused on identifying a possible “population structure” for reaction norms, that is, to summarize the impact of the historical plasticity patterns shared by the genotypes, which can be used as an additional genetic relationship matrix (GRM) for multi-environment predictions. The second is a new way to implement a weighted environmental relationship matrix (ERM), which is capable of accommodating past and future environmental covariables for predicting the expected “phenotypic similarity” for a given trait. Then, for the latter, different weights of each environmental factor should be achieved for different traits of the same genotype – which means that we can directly relate the quality of the growing conditions with the trait expression, translating the theoretical putative environmental influence to actual impact on the trait performance of the genotypes. Finally, a third “G× E pedigree” is born by combining the previous approaches, which result in a single-step G× E kernel considering reaction-norms (genotype-specific coefficients), genetic relationships (from pedigrees or molecular markers), field-trial (or location)-specific weights for each environmental factor and observed covariables in past field trials. This last is a more parsimonious manner of integrating different data sources capable of explaining the G× E variation for a given trait, at a given experimental network or breeding region. For this last application, it seems that a successful approach that follows a similar philosophy is the use of self-organizing maps (SOM) to identify genotype-specific responses for key environmental types combining historical yield and envirotyping data (Bustos-Korts et al., 2022).

Here, it was demonstrated that the conventional multi-environment GBLUP (M01, no enviromics) has poor resolution in explaining genotype-specific G× E variations for future years. As this models only accounts for genotypic covariates (G-matrix), and not any other environmental information, this result is in fact not a surprise. Conversely, our results suggest that the approaches (M04, M05 and M06) are a cost-effective and biologically accurate ways to correct this issue, with the benefit of better explore the cross-TPE and cross-season phenotypic data for training the GP models. Furthermore, the models M04-M06 aims to capture the historical background of seasonality variations by modeling the plasticity (reaction-norm matrix) of relatives while measures the actual weights of the envirotyping information in the phenotypic variation. Thus, this is an evolution of the conventional way to use raw environmental data to shape the similarity among locations.

A key point of our study was the development of the EPA-weighted location similarity matrix (***Φ***), used by itself (M04) or combined with other enviromic-aided approaches. The ERM derived from this matrix is modeled by nonlinear GK, bringing all the benefits of this approach, as already discussed in the last section. Perhaps an interesting point of this matrix is that it can always be updated (every year) and can also include new testing sites. For example, if we don’t have any phenotypic data for a new location (that is, we don’t have matrix ***Φ***_0_ considering this new location, Eq. 11), then we can still use the long-term envirotyping data for a new *j* location, assuming the weights for the *m* environmental covariables of this location 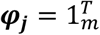, while the observed *r* location carries its specific weights computed by Eq. 11 using the data of the past experimental network; then ***φ*** = [***φ***_***r***_, ***φ***_***j***_]. This is a feasible and practical approach capable of using the outputs of the EPA analysis to unlock the avenue for predicting the expected phenotypic correlation for a new testing location. Finally, here we used a Hierarchical Bayesian approach, and the results demonstrated that the use of the kernels (***K***_***G***_, ***K***_***E***_ and ***K***_***GE***_) can be easily adapted for other computational platforms, such as convolutional neural networks (CNN) or assembly methods such as gradient boosting machine. Thus, a generic representation of a multi-environment genomic prediction could be given by ***y*** = *f*(***K***_***G***_, ***K***_***E***_, ***K***_***GE***_) + ***ε***, in other modeling forms, just considering markers (SNP) and supervised EPA-based outcomes, such as: ***y*** = *f*(*SNP*, ***φ, Λ***) + ***ε***.

### Environmental-Phenotype Association act as a reinforcement algorithm in recycling information from historical trialing networks

Finally, we can suggest a pipeline that combines our results with other approaches. For example, from the same multi-environment trial data we can follow two pathways. The first pathway is the “genotype analysis” based on the process of PLS 1 (univariate PLS for modeling the reaction-norms). This process has led to different purposes, but mostly to the estimation of the genotype-specific coefficients that can be part of cultivar targeting (e.g., Porker et al., 2020; Costa-Neto et al., 2020a), which could also be used as a strategy for reducing the dimensionality in genome-wide studies across multiple environments. For the latter, the use of the “reaction-norm coefficients” as a “trait” for genome-wide association studies (Li et al., 2021; Mu et al., 2022) cold help to understand the interplays between genomics and phenotypic plasticity. Here we introduce the reaction-norm matrix (**R** matrix) as containing “markers” of the plasticity patterns observed in past evaluated genotypes. Thus, instead of using a large data base of phenotypes from genotype x environment combinations, we can add the same information by recycling the past reaction-norms into a intuitive and synthetic matrix of “plasticity markers” (phenotype ∼environment variation associations).

The second pathway is the “environmental analysis”, which is based on the PLS 2 algorithm (multivariate PLS for modeling environmental weights). This approach is useful in creating more accurate ERM, which could be used to improve the accuracy of GP but also for identifying key environments and optimizing multi-environment trial efforts, which consequently could also extend the range of applications of the predictive analytic pipelines due to the possibility of better predicting the quality of some future environment. This last application can be used, for example, not only in the first multi-environment trial evaluations in breeding pipelines, but also in post-breeding stages aimed to support the decision about which locations are more likely to be successful as seed production fields, which is a key logistic aspect for multiplying seeds of the candidate cultivars.

Nonetheless, it is important to highlight two key aspects. First, every reaction-norm is just a sample of the possible reaction-norms that a certain genotype could experience due to its potential genetic-defined plasticity (Costa-Neto et al., 2021c). That is, a “snapshot” of its potential phenotypic plasticity as this reaction-norm coefficient is dependent on the diversity level of the MET and how it can represent the actual TPE. Because of this, some GP/GWAS models accounting for reaction-norms could carry noise related to the limits of the environmental diversity which each genotype experienced in the past. Thus, in the step of reaction-norm computation, the use of large historical data sets is key to reduce this possible bias. Secondly, but not less important, to work as reinforcement learning, this approach should be conducted after every year of trialing, where each new EPA is inserted into the data base and reaction-norms/environmental weights are fine-tuned.

## CONCLUSIONS

In this study we proposed a novel framework for the prediction of future seasons or new locations that parsimoniously integrates high-throughput environmental data (enviromics) and maker data to improve the prediction accuracy of a completely new environment (year or location). This new framework was tested using wheat multi-year data of CIMMYT and outperformed conventional multi-environment GBLUP methods by large margins because G×E signal was better accounted for and the noise in the environmental and marker data used as inputs was better removed. To be able to capture the essential patterns of the signal, the key was the partial least square algorithm that here was used in the first step as a way for variable selection of the marker and environmental data; then, in a second step, the selected new variables were used for training the multi-environment GBLUP models with which the predictions of unseen year or environments were finally performed. We call this approach that uses a partial least square algorithm for variable selection “Environment-Phenotype Associations (EPA)” since it allows capturing the essentials of G×E information from historical breeding data, to be able to use this refined information in the prediction models. Additionally, our framework allows capturing linear and nonlinear patterns since it was implemented under linear and nonlinear kernels, and allows identifying which genotypes and environments are the most important since it was possible to compute a Variable Importance in Projection (VIP) score. Finally, we encourage more empirical evaluations using our proposed framework to provide more empirical information about its advantages and disadvantages.

## Appendix

PLS regression is a technique used to combine the benefits of conventional ordinary least squares (OLS) and principal component analysis (Aastveit and Martens, 1986). Thus, it is useful for both predictive and exploratory analyses. Another important use of PLS analyses relies on the integration of environmental information in systems biology (Teixeira et al., 2011) and G× E analysis, this latter as an alternative to deal with the higher and natural collinearity of environmental factors and explain possible environmental causes of adaptation (Vargas et al., 1999; Monteverde et al., 2019; Porker et al., 2020). In fact, the first mention of “enviromics” did use this technique to support “reverse engineering” of the genomic features related to the dynamics within the cell environment (Teixeira et al., 2011). Due to its popularity and simplicity, we chose this approach to perform the so-called Envirotype-Phenotype Association (EPA) in two ways: (PLS 1) to estimate “adaptability descriptors” of the genotypes, that is, genotype-specific coefficients for each genotype and environmental factor; (PLS 2) to extract weights of environmental diversity, that is, associating environmental factors and the observed phenotypic-based similarity between environments. For both purposes, a generic representation of the PLS model is given below and its particular use in supporting the development of “enviromic-aided pedigrees” will be discussed in the next sections.

In the PLS algorithm, a matrix of predictors (**X**, *q* environments × *m* variables) is decomposed into three matrices: (1) a matrix of scores (**T**) (referred to as X-score), (2) a matrix of loadings (***P***’, X-loading, *m* × *a*) and (3) the residuals of this decomposition (**C**); thus:

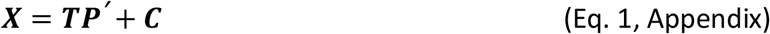

Simultaneously, the matrix of responses (***Y***) is decomposed into the three matrices of Y-scores (**U** matrix), Y-loadings (**<u>Q</u>***′* matrix), and the Y-residuals (denoted here as ***F***); thus:

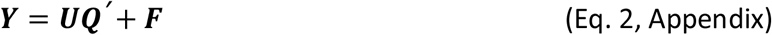

To estimate adaptability descriptors (PLS 1), we adopted the univariate PLS approach, considering the vector of phenotypes for each genotype (**Y** assumed as a vector of environmental-centered values for each genotype at each *q* environment × 1) of all observations across all test environments. In a certain way, this is a site-regression approach (SREG) (Crossa and Cornelius, 1997), where the main genetic effect plus genotype-specific G×E variation (G+GE) were attributed to the **y** vector. Here we are interested in modeling the combination of broad genetic effect (model intercept) plus a GE in terms of reaction-norms (Costa-Neto et al., 2020). To implement this, each model accounts for an independent single-genotype variation analysis over a matrix of environmental descriptors (***W***-matrix for environments, with *q* environments × k covariables). More details are given in the description of M05 in Genomics Prediction section.

On the other hand, for extracting the weights of environmental diversity, we considered the multivariate PLS approach (**Y** as a matrix, with dimension *q* × *q*, here *q* = number of locations), associating the long-term environmental descriptors for a given location (here a W-matrix for locations, with *q* locations × k covariables), and a matrix of phenotypic-based environmental similarity (S0-matrix) for each location. More details are given in the description of M05 in Genomics Prediction section.

In summary, the PLS algorithm aims to minimize the observed norm of ***F***, while looking to maximize the correlation between ***X*** and ***Y*** by the inner relation ***U***=***TD***, where ***D*** is a diagonal matrix. Consequently, the X-scores are orthogonal, assuring the control of collinearity among the original ***X*** predictors. This process results in the estimation of linear combinations of the original variables, computed from a matrix of weights (***L***), with a dimension of *a* × *b*, where *a* is the number of rows in the ***X*** matrix and *b* is the number of components (latent vectors) considered, with ***T***=***XL***. Finally, the linear coefficients to relate **X** and **Y** (referred here as **B**) are computed by using the iterative relation:

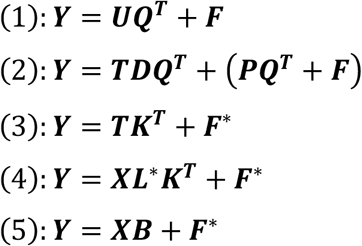

Thus, in summary,

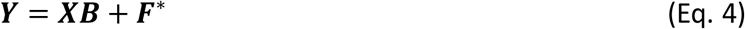

where the matrix of coefficients is computed by ***B*** = ***LK***^***T***^, and ***F***^*^ is the residual variance in **Y** not captured by the core of latent vectors (hereafter called LV). Here we adopted the nonlinear iterative partial least squares (NIPALS) algorithm to sequentially extract the PLS components. Detail on the NIPALS algorithm can be found in Palermo et al. (2009) and Sanchez (2012).

